# Precision fMRI reveals that the language network exhibits adult-like left-hemispheric lateralization by 4 years of age

**DOI:** 10.1101/2024.05.15.594172

**Authors:** Ola Ozernov-Palchik, Amanda M. O’Brien, Elizabeth Jiachen Lee, Hilary Richardson, Rachel Romeo, Moshe Poliak, Benjamin Lipkin, Hannah Small, Jimmy Capella, Alfonso Nieto-Castañón, Rebecca Saxe, John D. E. Gabrieli, Evelina Fedorenko

## Abstract

In adults, left hemisphere (LH) damage often leads to aphasia, but many cases of early damage leave linguistic processing intact, with a functional language system developing in the right hemisphere. To explain this early apparent equipotentiality of the two hemispheres for language, some have proposed that the language system is more bilateral during early development and becomes increasingly left-lateralized with age. We examined language lateralization using fMRI in two large developmental cohorts (total n=273 children aged 4-16 years; n=107 adults). Strong, adult-like LH lateralization (in response magnitude and activation volume) was evident by age 4, although other features of the LH language network showed protracted development, including the magnitude of language response and the strength of functional connectivity. Thus, although the RH can take over language function in some cases of early brain damage, this plasticity occurs in spite of adult-level LH bias present by age 4 years.

## Introduction

In a truly incredible feat, approximately six months after they are born, human babies begin to recognize words for common objects^1^, and toward their first birthday, they utter their first words^2^. Over the following couple of years, their vocabularies explode and they learn to combine words into phrases and sentences. Although linguistic abilities continue to increase in their sophistication into adolescence and even adulthood^3–5^, typically developing children can already understand and express complex ideas through language by 4 years of age^6^. But is the brain infrastructure for language in these young but already quite competent language users the same as in adults?

In adult brains, language processing, including both comprehension and production, draws on a specialized network of frontal and temporal areas^7,8^. In the vast majority of individuals, this network is lateralized to the left hemisphere (LH), which manifests as a) stronger and more spatially extensive responses to language in the left hemisphere (see^9^ for data from >800 individuals), and b) a greater likelihood of linguistic deficits (aphasia) following LH damage in adulthood^10^. One important controversy about the development of language-processing mechanisms concerns the degree of LH lateralization of the language network in children.

According to one influential proposal, the language system starts out more bilateral and becomes increasingly left-lateralized with age^11^. This proposal was put forward to explain the fact that LH damage at or shortly after birth and in early childhood can leave linguistic functions intact, with the RH homotopic areas taking over^12–16^ (see^17^ for a review). This proposal also makes predictions about language development in typical brains, which can be evaluated using functional brain imaging. In particular, the fronto-temporal language network should be more similar between the two hemispheres in its responses to language during early development compared to later childhood and adulthood. Indeed, some neuroimaging studies that have examined responses to language from about age 4 years onwards have reported more bilateral responses at younger ages and more strongly left-lateralized responses later in life^18–25^. However, other studies have reported no difference between children and adults in the degree of LH-lateralization of language responses^26,27^, including for children as young as 4 years of age^28^.

The complexity of the empirical landscape with respect to language lateralization in childhood may have to do with i) the predominant reliance of past fMRI studies on the traditional group-averaging approach, which suffers from low sensitivity, low functional resolution, and low interpretability^29–31^, and ii) the diversity of experimental paradigms, which complicates the across-study comparisons. Some paradigms further conflate language processing with other, more bilateral functions, including lower-level speech perception and general task demands. Speech perception draws on bilateral auditory areas, distinct from the amodal language areas^32,33^, and executive processes related to task demands (such as attention and working memory) recruit a bilateral fronto-parietal network—the Multiple Demand network^34,35^, which is also distinct from the language areas^36^, including in children^28^. For example, many language experiments require not only comprehension of words or sentences but also meta-linguistic judgments or comprehension questions, which are known to recruit the Multiple Demand network^37^. If these task components are more difficult for children than adults—given the protracted development of executive functions^38,4^—then language paradigms that include extra-linguistic task components may recruit the bilateral Multiple Demand network to a greater degree in children, which would lead to the appearance of more bilateral responses (we return to this issue in the <u>Discussion</u>).

Here, we characterize age-related changes in the language network across two independent developmental cohorts (Dataset 1: 206 children, aged 4-14 years, and 91 adults; Dataset 2: 67 children, aged 4-16 years, and 16 adults). We use a robust individual-subject fMRI approach (‘precision fMRI;’ ^29,39,40^) to account for inter-individual differences in the precise locations of language areas (see **SI-2B** for evidence of inter-individual topographic variability in our child participants). We also use an extensively validated language ‘localizer’ paradigm^30,41,9^, which isolates the language areas from both lower-level speech areas and domain-general areas sensitive to task demands^36,8^. Although our main focus is on lateralization, we additionally examine other properties of the language network, for which the developmental trajectory has also been debated in the developmental neuroscience literature. These include the LH language network’s topography (in particular, the emergence of the frontal-lobe component^22,42,24,43,44^, its magnitude of response to language, and the degree of functional connectivity among its regions^42,45^.

To foreshadow the results, across both datasets, we found that LH lateralization, based on either volume or magnitude of activation, is already adult-like by 4 years of age (see **SI-5** for generalization to a third, previously published dataset^25^). In contrast to the lateralization of language responses, the magnitude of response to language and the strength of inter-regional functional correlations in the dominant (left) hemisphere showed a pronounced increase from early to middle to late childhood, at which point they reached adult levels. The lateralization findings suggest that the capacity of the RH for language processing in cases of childhood LH damage does not depend on the language system being more bilateral in young children: at least, by age 4 years, an adult-like LH bias is already present. The magnitude and functional correlation results establish the normative developmental trajectory for these key individual-level neural markers of language, thus laying the groundwork for future investigations of linguistic processing at earlier ages and in developmental populations with communication disorders.

## Results

All participants (n=273 children aged 4-16 years; n=107 adults; <u>Methods-Section1</u>) performed an extensively validated language localizer task^30^ based on a contrast between listening to short age appropriate stories/passages (the critical condition) and a perceptually similar condition where linguistic content is not comprehensible (the control condition) (**Figure 1**; see Methods-Section2 and **SI-1A** for details). This localizer has been shown to engage the language areas—which support lexical access, syntactic structure building, and semantic composition^46^—and not to engage other nearby functional areas, such as the lower-level speech perception areas^33^ or areas of the domain-general Multiple Demand network^47^. All statistical analyses were performed on the brain measures extracted from the language areas, which were individually defined using the language localizer contrast (see <u>Methods-Section6,7</u>), including critically, measures of language lateralization based on the volume of activation and the magnitude of neural response.

**Figure 1.**
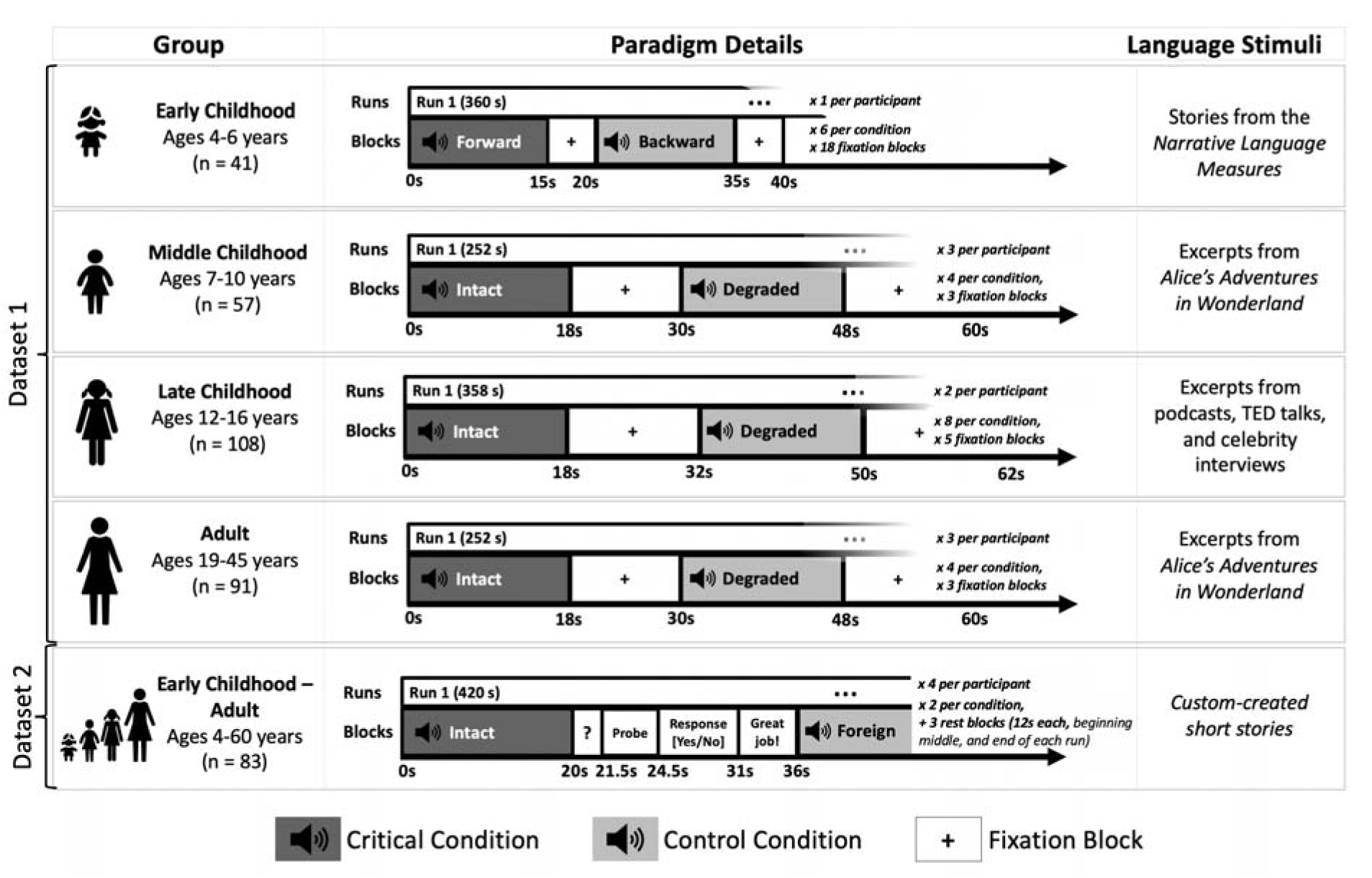
A summary of the design and procedure for the language localizer variants used for participants in Datasets 1 and 2. (see <u>Methods-Section2</u> and **SI-1A** for details). In Dataset 1, the paradigm was identical between the middle childhood and adult groups. In Dataset 2, the paradigm was identical across age groups. Importantly, the language localizer has been previously established to be robust to variation in the materials, task, modality of presentation, specific language, and the nature of the control condition^30,48,41^, which means that the data from these experiments can be straightforwardly combined and compared.

### 1. By 4 years of age, the left hemisphere language network shows adult-like functional topography

Because of past claims that the frontal-lobe component of the language network exhibits protracted development^43^, we first examined the general topography of the language responses across our different age groups, to decide which component(s) to focus on for the critical analyses of language lateralization. As shown in **Table 1** and **Figure 2A-B**, reliable responses to language processing are robustly present in both the temporal-lobe and frontal-lobe components of the LH network in children, including in the youngest developmental group examined here (4-6 year-olds).

**Figure 2.**
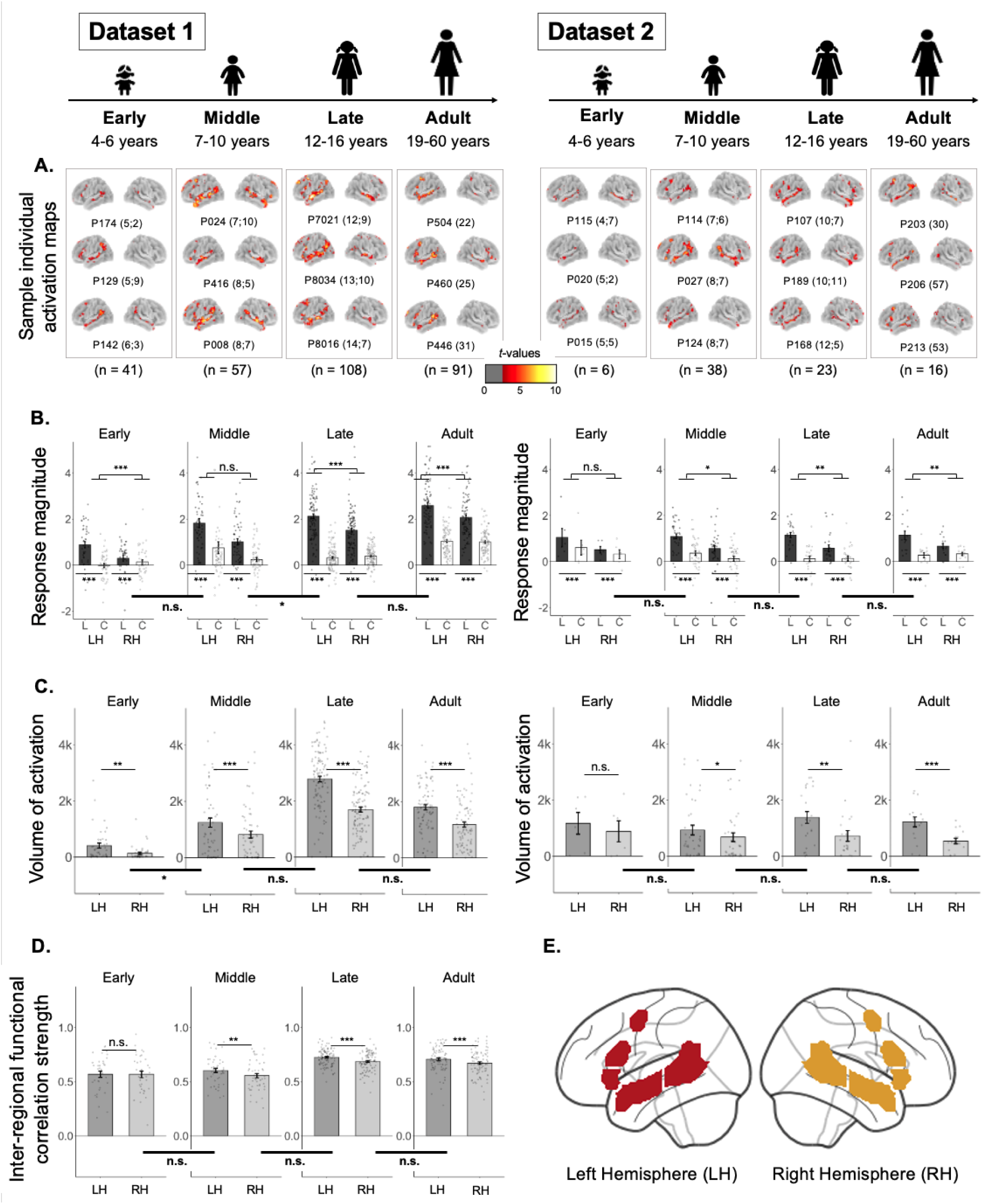
Responses to language and the strength of inter-regional functional correlations in the left hemisphere (LH) and right hemisphere (RH) language network in each age group. **A.** Sample individual whole-brain activation maps for the Language>Control contrast for participants in Datasets 1 and 2 (two broad columns) in each age group (four columns for each dataset). All maps are thresholded at the uncorrected whole-brain level of *p* < 0.01. For each age group in each dataset, we show three sample participants (all individual activation maps are available at https://osf.io/3mvpx/). Participants’ ages are shown in parentheses next to the participants’ unique identifiers (e.g., the age of participant P174 for the early childhood group in Dataset 1 is 5 years and 2 months). (These maps are used for visualization only; all statistical analyses are performed on neural measures extracted from these maps, as described in <u>Methods-Section7</u>.) **B.** Magnitude of response (in % BOLD signal change) to the Language condition (“L”, black) and the Control condition (“C”, white with black outline) relative to the fixation baseline (zero) in the LH and RH language network. Significant Language>Control effects are marked with asterisks below the x-axis (**Table 1**). Here and in C-D, significant effects of hemisphere are marked for each age group with asterisks above the bars in each plot (**Table 2**); and significant differences in the effects of hemisphere between age groups are marked on horizontal thick lines below the x-axis that straddle adjacent pairs of bar graphs (**Table 3**). **C.** Volume of activation (number of significant voxels; threshold: p<0.01) for the Language>Control contrast in the LH (dark gray) and RH (light gray). **D.** Strength of inter-regional functional correlation (Pearson’s moment correlation) among the language regions in Dataset 1 in the LH (dark gray) and RH (light gray) (no resting state data were collected for Dataset 2). **E.** The parcels that were used for fROI definition. These parcels (LH=red, RH=gold) were used to define the individual fROIs (for the measures in B and D; see **SI-2C** for visualizations of language fROIs in sample participants) or to constrain the voxel counts (for the measure in C) (see <u>Methods-</u> <u>Section7</u> for details). For all bars, dots correspond to individual participants, and error bars indicate standard errors of the mean by participant. Significance: ***=*p*<0.001; **=*p*<0.01; *=*p*<0.05; n.s.=not significant (see **Tables 1, 2 and 3** for details).

**Table 1.** Magnitude of response to language in the left hemisphere (LH) language network as a whole, and in the temporal and frontal components separately, across development. For each age group in each dataset (early childhood [Dataset 1 (DS 1) n=41; Dataset 2 (DS 2) n=6], middle childhood [DS 1 n=57; DS 2 n=38], late childhood [DS 1 n=108; DS 2 n=23], and adulthood [DS 1 n=91; DS 2 n=16]), we fit a linear mixed-effects regression model predicting the magnitude of the BOLD response (estimated in each region as described in <u>Methods-Section7</u>) from condition (Language vs. Control) with random intercepts for participants and fROIs (<u>Methods-Section8</u>). The responses are estimated in data independent from the data used to define the language fROIs. The *p-*values for the temporal and frontal areas are uncorrected, but all survive a Bonferroni correction for two comparisons.

| **Magnitude of response to the Language>Control contrast in the LH language network** |  |  |  |  |  |  |  |  |  |  |  |  |  |
| --- | --- | --- | --- | --- | --- | --- | --- | --- | --- | --- | --- | --- | --- |
|  |  | Language network |  |  |  | Temporal language areas |  |  |  | Frontal language areas |  |  |  |
|  |  | *b* | *SE* | *t* | *p* | *b* | *SE* | *t* | *p* | *b* | *SE* | *t* | *p* |
| DS 1 | Early | 0.91 | 0.08 | 11.2 | < **0.001** | 1.11 | 0.11 | 9.94 | < **0.001** | 0.77 | 0.11 | 7.06 | < **0.001** |
|  | Middle | 1.05 | 0.13 | 8.42 | < **0.001** | 1.55 | 0.19 | 7.96 | < **0.001** | 0.72 | 0.13 | 5.72 | < **0.001** |
|  | Late | 1.81 | 0.06 | 32.85 | < **0.001** | 2.53 | 0.079 | 32.15 | < **0.001** | 1.32 | 0.07 | 20.32 | < **0.001** |
|  | Adult | 1.54 | 0.06 | 24.88 | < **0.001** | 1.78 | 0.11 | 24.86 | < **0.001** | 1.34 | 0.07 | 18.2 | < **0.001** |
| DS 2 | Early | 0.44 | 0.1 | 4.58 | < **0.001** | 0.56 | 0.16 | 3.41 | **0.01** | 0.35 | 0.1 | 3.54 | < **0.01** |
|  | Middle | 0.73 | 0.06 | 11.75 | < **0.001** | 0.84 | 0.09 | 9.2 | < **0.001** | 0.66 | 0.08 | 8.02 | < **0.001** |
|  | Late | 1.03 | 0.07 | 14.34 | < **0.001** | 1.1 | 0.09 | 11.78 | < **0.001** | 0.97 | 0.1 | 10.05 | < **0.001** |
|  | Adult | 0.86 | 0.1 | 9.04 | < **0.001** | 0.81 | 0.11 | 7.45 | < **0.001** | 0.9 | 0.14 | 6.53 | < **0.001** |

**Table 2.** A LH bias in response to language in the language network across development. A. Effect of hemisphere on the magnitude of the Language>Control contrast. For each age group in each dataset, we fit a linear mixed-effects regression model predicting the average effect size (averaged across fROIs; effects sizes were estimated as described in <u>Methods-Section7</u>) from hemisphere with random intercepts for participants (including random intercepts for fROIs led to the model not converging). **B. Effect of hemisphere on the volume of activation for the Language>Control contrast.** For each age group in each dataset, we fit a linear mixed-effects regression model predicting the total number of voxels (across fROIs; number of voxels was calculated as described in <u>Methods-Section7</u>) from hemisphere with random intercepts for participants.

| A. Effect of hemisphere on the magnitude of the Language>Control contrast |  |  |  |  |  |
| --- | --- | --- | --- | --- | --- |
|  |  | *b* | *SE* | *t* | *p* |
| DS 1 | Early | 0.75 | 0.17 | 4.29 | < 0.001 |
|  | Middle | 0.30 | 0.15 | 2.04 | 0.046 |
|  | Late | 0.69 | 0.08 | 8.43 | < 0.001 |
|  | Adult | 0.48 | 0.07 | 6.66 | < 0.001 |
| DS 2 | Early | 0.24 | 0.19 | 1.29 | 0.26 |
|  | Middle | 0.29 | 0.08 | 3.81 | 0.001 |
|  | Late | 0.59 | 0.15 | 3.96 | **0.001** |
|  | Adult | 0.52 | 0.11 | 4.88 | **< 0.000** |
| **B. Effect of hemisphere on the volume of activation for the Language>Control contrast** |  |  |  |  |  |
|  |  | *b* | *SE* | *t* | *p* |
| DS 1 | Early | 272.2 | 88.83 | 3.06 | **0.004** |
|  | Middle | 425.21 | 90.4 | 4.7 | **< 0.001** |
|  | Late | 1087.91 | 87.3 | 12.46 | **< 0.001** |
|  | Adult | 614.09 | 79.32 | 7.74 | **< 0.001** |
| DS 2 | Early | 285 | 202.84 | 1.41 | 0.22 |
|  | Middle | 250.58 | 90.29 | 2.78 | **0.01** |
|  | Late | 656.65 | 185.64 | 3.54 | **0.002** |
|  | Adult | 680.9 | 119.9 | 5.68 | **<0.001** |

**Table 3.**
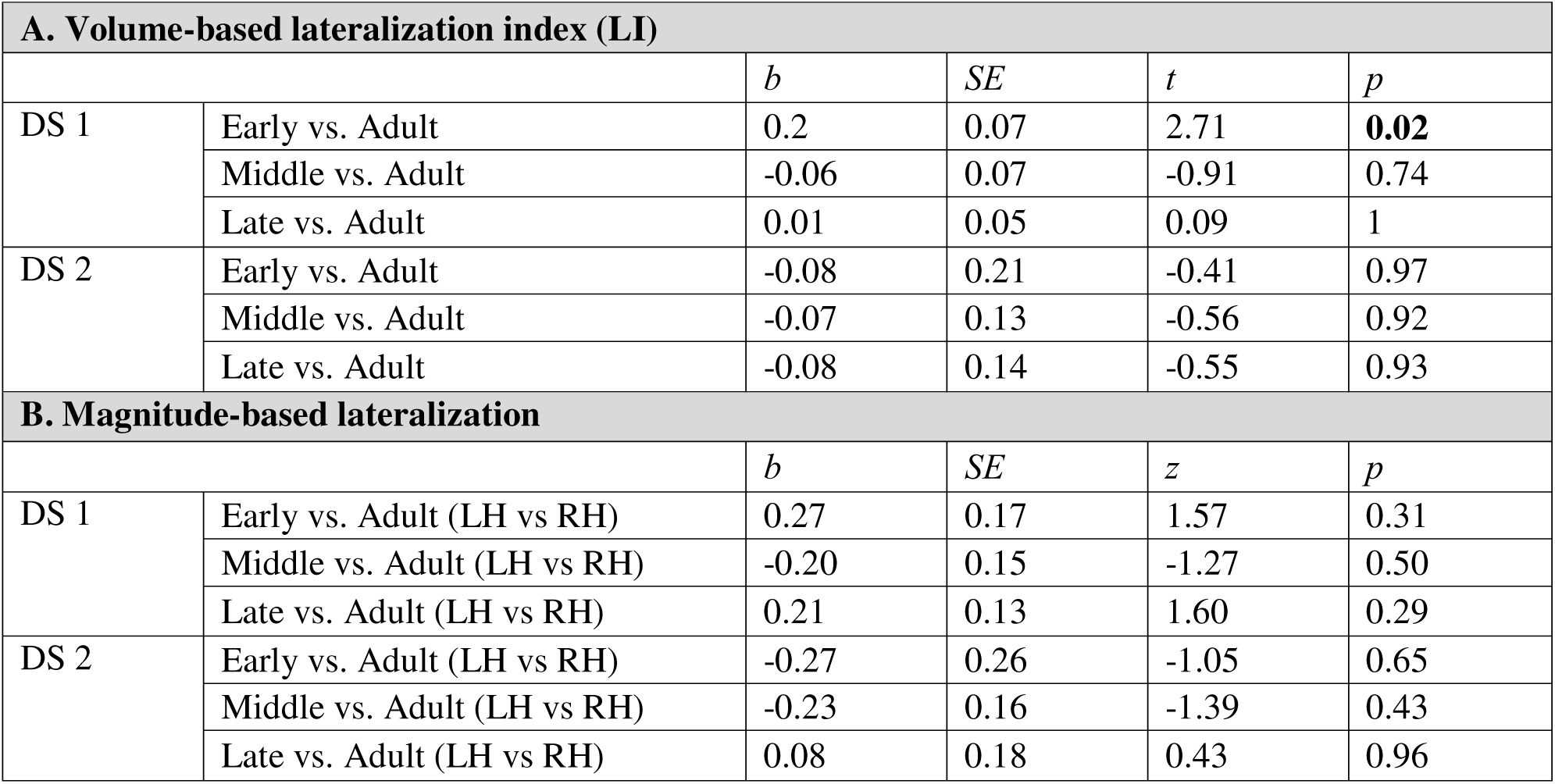
No evidence for an age-related increase in the LH bias for language. **A. Volume-based LI analysis.** We fit a linear regression model predicting LI (averaged across fROIs) from age group, followed by pairwise comparisons between the adult group and each child group, adjusted using Sidak’s method (<u>Methods-</u> <u>Section8</u>). (For the analysis that uses ordered age comparisons, see **SI-3F**; for the analysis that treats age as a continuous variable, see **SI-4B**.) (Note that the only significant effect observed here goes in the opposite direction of the one predicted by the hypothesis we are evaluating^11^.) **B. Magnitude-based lateralization analysis.** We fit a linear mixed-effect regression model predicting the size of the Language>Control contrast from age group, hemisphere, and their interaction, with random intercepts and slopes for hemisphere within participants and fROIs, followed by pairwise comparisons, as in A.

| **A. Volume-based lateralization index (LI)** |  |  |  |  |  |
| --- | --- | --- | --- | --- | --- |
|  |  | *b* | *SE* | *t* | *p* |
| DS 1 | Early vs. Adult | 0.2 | 0.07 | 2.71 | **0.02** |
|  | Middle vs. Adult | -0.06 | 0.07 | -0.91 | 0.74 |
|  | Late vs. Adult | 0.01 | 0.05 | 0.09 | 1 |
| DS 2 | Early vs. Adult | -0.08 | 0.21 | -0.41 | 0.97 |
|  | Middle vs. Adult | -0.07 | 0.13 | -0.56 | 0.92 |
|  | Late vs. Adult | -0.08 | 0.14 | -0.55 | 0.93 |
| **B. Magnitude-based lateralization** |  |  |  |  |  |
|  |  | *b* | *SE* | *z* | *p* |
| DS 1 | Early vs. Adult (LH vs RH) | 0.27 | 0.17 | 1.57 | 0.31 |
|  | Middle vs. Adult (LH vs RH) | -0.20 | 0.15 | -1.27 | 0.50 |
|  | Late vs. Adult (LH vs RH) | 0.21 | 0.13 | 1.60 | 0.29 |
| DS 2 | Early vs. Adult (LH vs RH) | -0.27 | 0.26 | -1.05 | 0.65 |
|  | Middle vs. Adult (LH vs RH) | -0.23 | 0.16 | -1.39 | 0.43 |
|  | Late vs. Adult (LH vs RH) | 0.08 | 0.18 | 0.43 | 0.96 |

In particular, consistent with what has been reported previously in adults (e.g., see^41^ for the results for the adult group in Dataset 1), participants in each developmental group showed a reliably stronger response during the Language condition compared to the Control condition (**Figure 2B**). This effect held across the LH language network as a whole in both datasets and every age group (all *p*s<0.001; **Table 1**). The effect also held for the temporal-lobe and frontal-lobe components of the network separately, again in both datasets and every age group (all *p*s<0.001, except the early group in Dataset 2 (n=6): *p*s≤0.01; **Table 1**). (For the parallel analyses of the homotopic RH language network, see **Figure 2A-B** and **SI-4A**.)

These results suggest that we can examine both the frontal-lobe and temporal-lobe components of the language network for our critical question of language lateralization across childhood, instead of restricting our analyses to the temporal-lobe component.

### 2. The language network exhibits an adult-like left-hemispheric bias by 4 years of age

In adults, the left-hemispheric (LH) bias for language manifests as a stronger (higher magnitude) and more spatially extensive (greater number of significant voxels) response in the LH language areas. We examined both of these measures in children, including in relation to adults.

First, we asked whether a LH bias was present in each developmental age group. The effect of hemisphere on the magnitude of the Language>Control effect held in all age groups in both datasets (*p*s<0.05; **Figure 2B**, **Table 2A**), except the early childhood group in Dataset 2. The effect of hemisphere on the volume of activation for the Language>Control contrast also held in all age groups in both datasets (*p*s<0.05; **Figure 2C**, **Table 2B**), except the early childhood group in Dataset 2. The lack of significant effects in the early childhood group in Dataset 2 is likely due to a small sample size of n=6 (note though that the mean inter-hemispheric differences are similar in size to the other groups; e.g., in the middle childhood group, there are, on average, 32% more significant Language>Control voxels in the LH than the RH, and in the early childhood group, there are, on average, 37% more voxels in the LH than the RH). For the groups that showed reliably lateralized responses at the network level, the effects also held for the temporal and frontal components of the network separately, for both the magnitude and the volume measures (all *ps*<0.05; **SI-3)**.

Next and critically, we asked whether the LH bias was larger in adults compared to the child groups. To do so, we used three complementary measures, in line with recent emphasis in the field on robustness to analytic choices^49–51^, and the lack of agreement on the best measure in laterality research^52,53^. In particular, we examined traditional lateralization index (LI) based on activation volumes^54,55,19^, lateralization based on response magnitudes^52,56^, and LI based on magnitudes that uses adaptive thresholding^57^. As detailed next, all three measures yielded the same answer of no evidence for an increase in lateralization between childhood and adulthood (**Figure 3**, **Table 3**).

**Figure 3.**
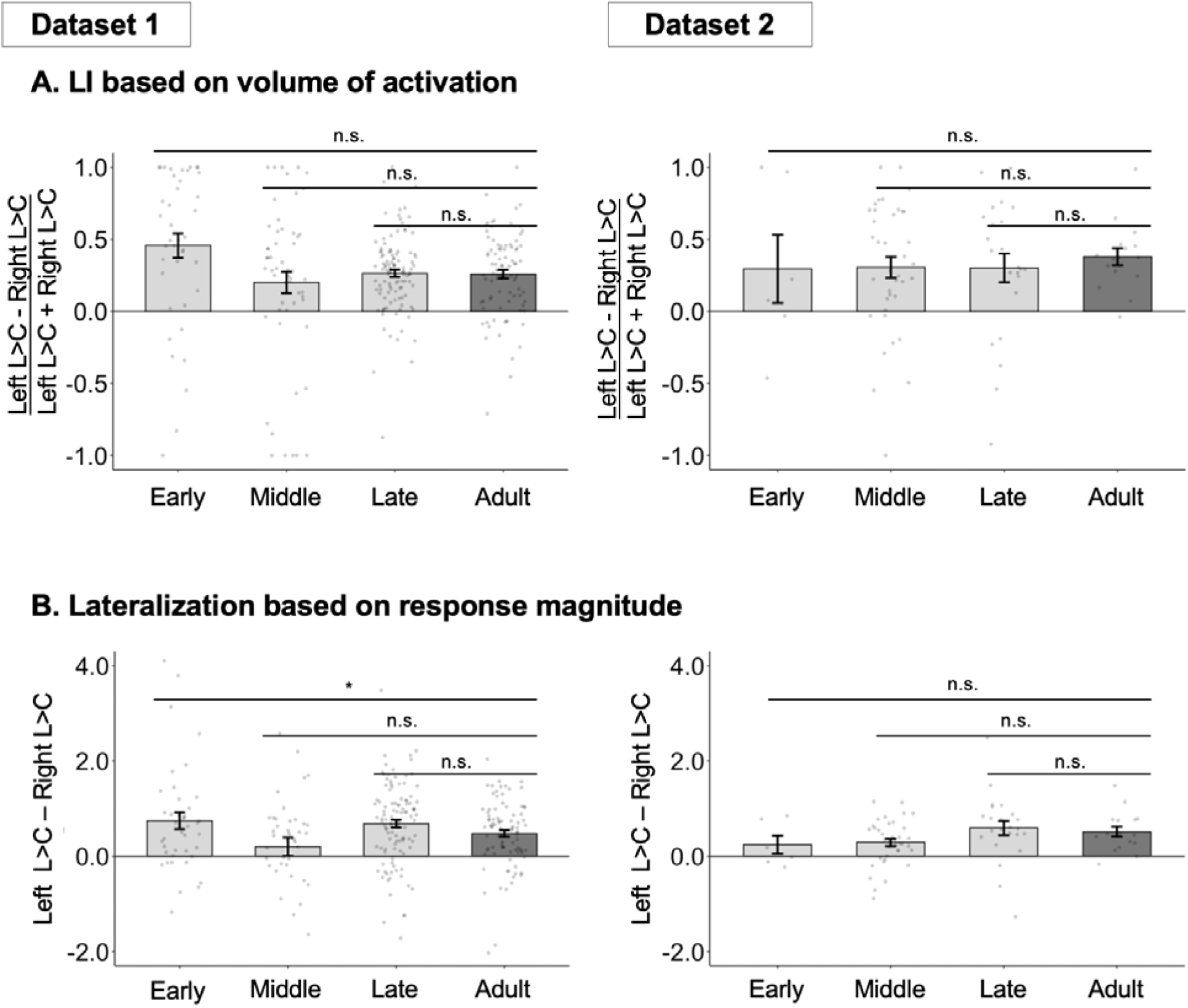
Developmental changes in lateralization. **A.** Volume-based LI measure for participants in Datasets 1 and 2 (two broad columns) in each child group (light grey bars) and the adult group (dark grey bar). Here and in B, significant differences between children and adults are marked for each child group above the bars (note that the only significant effect goes in the opposite direction of the one predicted by the hypothesis we are evaluating(Lenneberg, 1967)). **B.** Magnitude-based lateralization measure. For all bars, dots correspond to individual participants, and error bars indicate standard errors of the mean by participant. Significance: ***=*p*<0.001; **=*p*<0.01; *=*p*<0.05; n.s.=not significant (see **Table 3** for details).

The traditional lateralization index (LI) is based on the number of significantly activated voxels per hemisphere (for the Language>Control contrast) using the following formula: (LH − RH) / (LH + RH), where LH and RH represent the number of significant voxels in the left and right hemispheres, respectively. We found no evidence for age-related increase in lateralization (**Figure 3A**, **Table 3A**; see **SI-3A** for the results of frontal and temporal areas separately; see **SI-4B** for analyses where age was treated as a continuous variable), although DS2 was small and likely underpowered (see **SI-3D** for analyses that combine the two datasets and still find no age-related increase in lateralization). The traditional LI measure controls for group differences in the overall extent of activation^58,19,59^ but may be affected by differences in activation magnitude and is sensitive to outliers^57^. We therefore additionally examined differences in response magnitude between the two hemispheres. We again found no evidence for age-related increase in lateralization (**Figure 3B**, **Table 3B**). In addition, Bayesian analyses of the magnitude-based lateralization measure provided substantial-to-strong evidence for the null hypothesis of no monotonic age effects on lateralization in Dataset 1, with inconclusive evidence in Dataset 2 likely due to the small sample size (**SI-3B**). Finally, we used the LI toolbox developed by Wilke and colleagues^60,57^, which allows for adaptive threshold selection to derive a threshold-independent LI with a confidence interval computed via bootstrapping. The results (reported in (**SI-3C**) converged with the other two approaches showing no evidence for age-related lateralization increase.

### 3. Language response magnitude and functional connectivity in the LH language network increase between ages 4 and 16 and reach maturity by late childhood

In addition to lateralization, we examined developmental changes in the magnitude of response to language and the strength of inter-regional functional correlations in the language network in the dominant, left hemisphere. Both of these properties showed gradual increase across childhood (**Figure 4**).

**Figure 4.**
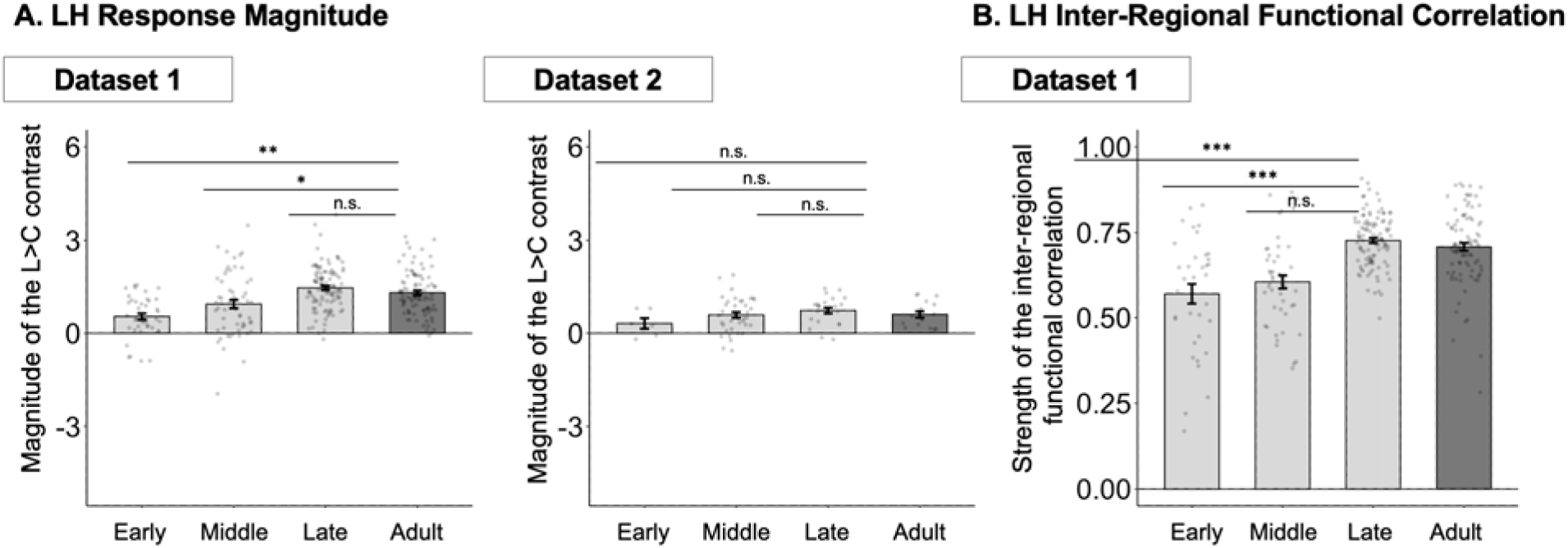
Developmental changes in language response magnitude and functional connectivity in the LH language network. **A.** Magnitude of the Language>Control contrast in Datasets 1 in each child group (light grey bars) and the adult group (dark grey bar); see the inset for Dataset 2 results. Here and in B, significant differences between children and adults are marked for each child group above the bars. **B.** Strength of inter-regional functional correlations in Dataset 1. For all bars, dots correspond to individual participants, and error bars indicate standard errors of the mean by participant. Significance: ***=*p*<0.001; **=*p*<0.01; *=*p*<0.05; n.s.=not significant (see **Table 4** for details).

**Table 4.** Evidence for age-related increases in language response magnitude and functional connectivity in the LH language network. **A. Magnitude of the Language>Control contrast analysis.** We fit a linear mixed-effect regression model predicting response magnitude of the LH language network from age group, with random intercepts for participants and fROIs. We then performed pairwise comparisons between the adult group and each child group, adjusted using Sidak’s method. (For the analysis that treats age as a continuous variable, see **SI-4B**.) **B. Inter-regional functional correlation strength analysis.** We fit a linear regression model predicting inter-regional correlations (averaged across the 10 fROI pairs for the LH network) from age group, followed by pairwise comparisons, as in A.

| **A. Magnitude of the Language>Control contrast in the LH language network** |  |  |  |  |  |
| --- | --- | --- | --- | --- | --- |
|  |  | *b* | *SE* | *t* | *p* |
| DS 1 | Early vs. Adult | -0.64 | 0.19 | -3.46 | **0.002** |
|  | Middle vs. Adult | -0.48 | 0.17 | -2.89 | **0.01** |
|  | Late vs. Adult | 0.26 | 0.14 | 1.87 | 0.17 |
| DS 2 | Early vs. Adult | -0.43 | 0.29 | -1.47 | 0.38 |
|  | Middle vs. Adult | -0.13 | 0.18 | -0.72 | 0.85 |
|  | Late vs. Adult | 0.16 | 0.2 | 0.83 | 0.8 |
| **B. Inter-regional functional correlation strength in the LH language network** |  |  |  |  |  |
|  |  | *b* | *SE* | *t* | *p* |
| DS 1 | Early vs. Adult | -0.14 | 0.02 | -6.14 | **< 0.001** |
|  | Middle vs. Adult | -0.1 | 0.02 | -5.08 | **< 0.001** |
|  | Late vs. Adult | 0.02 | 2 | 1.16 | 0.57 |

In Dataset 1, the Language>Control effect was larger in the adult group compared to the early and middle childhood groups (*ps*<0.05; **Figure 4A-left, Table 4A;** see **SI-4C** for evidence that the results are similar when controlling for head motion; see **SI-4B** for analyses where age was treated as a continuous variable), but not compared to the late childhood group. In Dataset 2, the qualitative pattern was similar—with responses increasing from early to middle to late childhood and not showing a further increase between late childhood and adulthood—but none of the pairwise comparisons reached significance (**Figure 4A-right, Table 4A**). The analysis where age was treated as a continuous variable revealed a positive effect of age on the magnitude of the Language>Control contrast in Dataset 1 (*p*<0.001; **SI-4B**); this effect was also positive albeit not significant in Dataset 2. The pattern of increasing response magnitude was also similar in the RH language network across both datasets (**SI-4A**).

The results for inter-regional functional correlations, as measured during a resting state scan, mirrored the results for response magnitude: the correlations were stronger in the adult group compared to the early and middle childhood groups (*p*s<0.001; **Table 4B**; see **SI-4C** for evidence that the results are similar when controlling for head motion^61^, but not compared to the late childhood group. The analysis where age was treated as a continuous variable revealed a positive effect of age (*p*<0.001; **SI-4B**).

## Discussion

We investigated age-related changes in the degree of language lateralization and in other properties of the fronto-temporal language network using two independent relatively large developmental fMRI datasets and an extensively validated language ‘localizer’ paradigm^30^. Using robust individual-subject analyses, which account for variability across individuals in the precise locations of functional areas, we found a strong and adult-like left hemisphere (LH) language bias even in our youngest child group (4-6 year-olds), with no evidence for an age-related increase in lateralization. In contrast, other features of the language network show a clear developmental change. In the remainder of the Discussion, we position these findings in the context of the prior literature and discuss their broader implications.

### By 4 years of age, language processing is as strongly left-lateralized as in adults

One important claim in the literature has been that the language network starts out as more bilateral and only becomes strongly lateralized to the left hemisphere with age^11^. This hypothesis was put forward to explain a difference in the likelihood of language deficits following damage to the LH in mature vs. child brains. In particular, LH damage in adulthood typically results in aphasia^10,62^. In stark contrast, damage to the LH in children can leave language ability largely preserved, including cases of damage occurring around birth^13,14,16^, in young children^13^, and even in some adolescents^63^ (cf.^64–66^). Functional imaging has revealed that in such cases, language processing is typically supported by the homotopic right-hemisphere frontal and temporal areas^12–16^ (see^17^ for a review). This ability of the right hemisphere to take on language function has led to arguments that RH frontal and temporal areas support language processing alongside the LH areas in developing brains, and the role of the RH areas gradually lessens with age, reducing its ability to take over language function^67^.

This hypothesis—whereby the language system starts out more bilateral—makes a testable prediction for typical development: the LH and RH language areas should respond more similarly to language in young children. Over the course of development, these responses are expected to gradually shift towards an adult-like profile where the LH areas show stronger and more extensive responses to language compared to the RH areas. Here, across two datasets and three measures of lateralization, we do not find support for this prediction: the language network is already left-lateralized in 4-6 year-olds (complementing some earlier studies; e.g.,^26–28^), and the degree of lateralization is similar to that in adults. In other words, we find no evidence for age-related increase in language lateralization from age 4 through adulthood a) across all standard lateralization measures, b) with Bayesian analyses providing substantial-to-strong evidence for the null hypothesis of no age-related increase in lateralization, and c) with robust evidence for age-related changes in other aspects of brain responses to language.

### Prior findings of age-related increases in lateralization may be due to conflating linguistic and task demands

Why have some prior studies^19–21,23,68,24,25^ found more bilateral responses to language in childhood? An important contributor may be the use of paradigms that conflate linguistic demands (i.e., lexical retrieval, syntactic structure building, and semantic composition)— supported by the language network (e.g., ^8^ —and general task demands (requiring attention, working memory, response selection, etc.). In particular, many language paradigms that require overt responses (e.g., the commonly used verb generation task^18,19,69^ tax both language processing and general executive abilities. The latter are supported by the domain-general Multiple Demand (MD) network^34,35^—a bilateral network of frontal and parietal areas whose left frontal component is adjacent to, but distinct from, the language areas^70,71,36,72^; (see^37^ for evidence that language paradigms accompanied by task demands recruit both the language and the MD network). Given that most tasks are more difficult for younger children, they may recruit the MD network to a greater extent. Because the MD network is bilateral, its greater recruitment will manifest as more bilateral responses at younger ages. In other words, what has been interpreted as an age-related increase in the degree of LH lateralization of the language network may instead reflect an age-related reduction in the reliance on the (bilateral) MD network, and increased reliance on the (left-lateralized) language network.

We believe this issue affects an influential study by Olulade and colleagues^25^. Olulade et al. examined responses to language in children (similar age range as in the current study) and adults, and found more bilateral activity in children, and increasingly left-lateralized responses with age. However, their paradigm included both linguistic and task demands, and the task was more difficult in the critical language condition (listening to sentences and judging their factual correctness) than in the control condition (listening to sentences played backwards and pressing a button when a beep is present at the end). The behavioral data further suggest that the critical task was more difficult for younger children. Besides, brain responses were measured in large anatomical masks and lateralization was computed using a newly introduced measure. When Olulade et al.’s data are analyzed in more restricted language-related regions of interest using standard lateralization measures, we find that even the youngest children (age 4-6 years) already show strongly left-lateralized responses, similar to the current study (see **SI-5C**).

Future work on the development of the language network should take care to separate the language network from the domain-general Multiple Demand network, and other networks known to be functionally distinct from the language network. One way to do this is to adopt paradigms that focus on passive comprehension, which minimize task demands (e.g., contrasting passive comprehension of language with perceptually matched control conditions^30,48^. If a task is included, care should be taken to a) ensure that the task in the control condition is at least as difficult as in the critical condition, and/or b) separate the task temporally from the language-processing component so that it can be modeled separately (as in Dataset 2 here), and/or c) use independent functional localizers or at least spatial priors from pre-existing large-scale datasets from validated localizers (e.g.,^9^).

### Reconciling early left -hemispheric language lateralization with early equipotentiality of the two hemispheres for language

Given that language processing is already lateralized to the left hemisphere by age 4 years, how do we square this finding with evidence from early LH brain damage, which often leaves language processing unimpaired? Existing evidence unequivocally shows that the two hemispheres *are* equipotential for language early in life (e.g.,^13–16^). Evidently, however, this equipotentiality does not manifest as similar responses to language in the two hemispheres. One possibility is that brain imaging studies, including ours, are measuring language responses too late in the developmental trajectory: maybe language processing is more, or even fully, bilateral until 1-3 years of age. Some evidence exists of LH lateralization prior to age 4^28^ and even prior to age 2(Olson et al., 2025) although children in those cohorts could still be too old. Further, at least some cases have been reported where LH damage occurs after age 4 years, but no long-term language deficits ensue^13,63^, but premorbid language lateralization in these individuals is not known.

Alternatively, an age-related change in the ability of the RH areas to take over language function may be caused not by the reduction in the RH areas’ responses to language (see **Figure 2** and **SI-4A** for evidence of consistent recruitment of the RH for language processing in adulthood; also, see^74,9^) but because the RH areas become increasingly specialized for processing non-linguistic information. For example, RH temporal and frontal areas in adults have been implicated in visual and auditory social perception^75–78^, mental state reasoning^79^ and processing prosodic information^80,81^. In contrast, the LH language areas remain language-selective^8^.

Yet another possibility is that other aspects of brain development mediate early equipotentiality. For example, age-related decreases have been reported in the degree of functional connectivity between the LH language areas and their RH homotopic areas^82^. Perhaps the ability for inter-hemispheric exchange of information is what allows the RH language areas to take on language function early during development but not later in life. However, in our data, we do not see a pattern of decreasing inter-hemispheric functional connectivity (**SI-3E**).

It is also worth noting that early equipotentiality of the two hemispheres for language— accompanied by bilateral responses to language—may only be present under the conditions of atypical brain development. Indeed, more bilateral responses to language have been reported in individuals with neurodevelopmental conditions, such as autism^83,84^, dyslexia^85^ and epilepsy^86^. If atypical brain development is a ‘prerequisite’ for more bilateral language processing —and thus easier reorganization of language function to the right hemisphere—we would expect easier/faster language recovery in children with epilepsy who undergo LH surgical resections compared to children who experience LH stroke. To our knowledge, no direct comparisons on matched populations have been performed to evaluate this prediction.

Finally, the brain’s remarkable capacity to reorganize in response to injury or atypical experience need not imply the lack of functional biases in typical brains. After all, even in adulthood— where strong LH language lateralization is not disputed—many individuals with stroke aphasia show complete or near-complete recovery^87^, although recovery mechanisms remain an active area of research^88^.

### Some features of the LH language network change across development and reach maturity by late childhood

Although linguistic abilities show a high degree of sophistication by age ∼4 years (e.g.,^89,90^, many of these abilities continue to develop well into the late teens^3–5^. In line with this protracted developmental trajectory, two aspects of the brain organization for language processing showed a developmental change. First, the strength of the response during language comprehension relative to a control condition increased from early, to middle, to late childhood, with no difference between the late-childhood and adult groups. This pattern was consistent between the two datasets, despite differences in the populations tested and details of the experimental materials and paradigm. Second, the strength of inter-regional functional correlations during unconstrained cognition (resting state) showed an increase across development (including after controlling for head motion; **SI-4C**), also reaching adult-like levels by late childhood.

Developmental changes in both the magnitude of response to language and in the strength of inter-regional functional correlations have been reported in past studies (e.g.,^22,42,24,43,44^. However, to the best of our knowledge, none of those studies have relied on paradigms that could differentiate language areas from areas of nearby functionally distinct networks (e.g.,^70,47,72^. Using a paradigm that selectively engages language areas and yields reliable estimates of brain activity at the individual level in the current study allowed us to unambiguously attribute the observed neural changes to the maturation of linguistic abilities and their underlying substrates (see also^28^).

A developmental increase in response magnitude to language may reflect relatively experience-independent biological maturation of the underlying neural circuits^91^ or experience-related changes in linguistic ability. One indirect source of evidence for the experiential account comes from responses to language in brains of adults learning a new language. In particular, Malik-Moraleda, Jouravlev et al.^92^ found that neural responses to language scale with proficiency levels. Thus, stronger responses in the language network may reflect better linguistic ability in both children learning their first language and adult second-language learners(see also ^93^ for behavioral evidence for experience-based accounts of language learning). However, it remains important to directly relate neural responses to language with behavioral measures of linguistic ability in young language learners, as well as to understand how changes in the ability to pay attention across development may affect those responses.

### Limitations and Open Questions

The current study is limited in several ways. First, we adopted a cross-sectional approach. Tracking changes in the brain infrastructure of language processing longitudinally may offer greater sensitivity in detecting age- and experience-related changes. Second, we did not attempt to match the age groups on any variables that may affect brain responses (e.g., motion or performance on language measures). This concern is somewhat ameliorated by the inclusion of two datasets and the observation of similar patterns across them; nevertheless, more carefully matched samples performing an identical version of the language paradigm should be examined in future research, and generalized to other paradigms^94^, including those that tax language production demands. Finally, all the developmental data come from English, which is not representative of the world’s languages (e.g.,^95^), leaving generalization to other languages an important future direction.

Several questions remain about how the language network develops. First, when and how do the language brain areas emerge? The youngest children in the current dataset are ∼4 years old. This is also true of most past studies of brain language development (e.g.,^19,21,96^. By age 4 years, children can understand and express complex ideas through language^89,90^. The field sorely lacks brain data from infants and toddlers (age range: 6 months to ∼3 years old), but probing brain responses to meaningful language during the first years of life using spatially precise brain-imaging methods like fMRI is extremely challenging(but see^73^).

Second, how does specialization for language processing develop? The language brain areas in adults are strongly specialized for language relative to diverse non-linguistic inputs and tasks^8^. In the current study, we only examined responses to language relative to one control condition (perceptually similar but not meaningful stimuli), thus leaving open the question of whether language areas perform additional functions in children. A richer characterization of these areas’ response profile across development remains an important future goal (see^28^ for evidence of selectivity of the language areas relative to a demanding working memory task in young children).

Third, what is special about the left hemisphere such that in most individuals, language processing ends up localized to the left? Hypotheses about the LH bias for language are plentiful^97,98^, but obtaining a clear answer has proven challenging. Understanding whether the language system is already lateralized when it first emerges (the first open question above) and whether or how selectivity for language changes across development (the second question) may provide some clues. A few studies have examined responses to speech in infants: some have argued for a LH bias^99,42^, but others found no evidence for lateralization^100,101^. However, speech perception is an earlier-emerging ability which is distinct from language comprehension^8^ and largely bilateral in adult brains^32,33^, so those studies do not directly inform the lateralization of the language system.

In conclusion, the language system of young children is as strongly lateralized to the left hemisphere as in adults, which challenges the hypothesis that the language system is more bilateral early in development^11,67^. Although the level of response to language and the inter-connectedness of the language network continue to increase into late childhood, adult-like lateralization to the left hemisphere is already present by age 4 years.

## Materials and Methods

### 1. Participants

All participants were recruited from the greater Boston area of the United States, via flyers and advertisements on social media. Researchers aimed to recruit half males and half females, based on self-reported sex assigned at birth, in each age group. Gender data were not collected in the present study. For all child participants, a parent or guardian provided written informed consent, and the child provided assent; adult participants provided written informed consent, in accordance with the Committee on the Use of Humans as Experimental Subjects at the Massachusetts Institute of Technology.

#### Inclusion criteria

Across datasets, child and adult participants were included in the analysis based on the following criteria: 1) being a native speaker of the language used in the experiment (English for all groups except for the adult group in Dataset 1; diverse languages for the adult group in Dataset 1, as described in^41^, 2) having no diagnosis of neurological disorder, and 3) having normal or corrected-to-normal vision. For Dataset 1, it was additionally ensured that child participants were born full-term (>37 weeks), had no history of brain injury and hospitalizations, and were not using psychotropic medications.

#### Dataset 1

After excluding 28 child participants and 0 adult participants from group-level analyses due to excessive motion (<u>Methods-Section5</u>; for summary of motion outliers by group, see https://osf.io/3mvpx/), the final Dataset 1 included 206 children (ages 4-14; average = 10.69, st. dev. = 3.18; 106 females; 178 right-handed) and 91 adults (ages 19-45, average = 27.40, st. dev. = 5.5; 48 females; 88 right-handed). The developmental data came from three projects that each focused on a different age range and did not concern the development of the language network (for details, see https://osf.io/3mvpx/): early childhood (n=41; ages 4-6, average = 5.78, st. dev. = 0.69), middle childhood (n=57; ages 7-10, average = 9.00, st. dev. = 0.74), and late childhood (n=108; ages 12-14, average = 13.45, st. dev. = 0.69).

#### Dataset 2

After excluding 31 child participants and 0 adult participants from group-level analyses due to excessive motion (<u>Methods-Section5</u>; for summary of motion outliers by group, see https://osf.io/3mvpx/), the final Dataset 2 included 67 children (ages 4-16; average = 11.5, st. dev. = 1.4; 29 females; 70 right-handed) and 16 adults (ages 18-60, average = 43.8, st. dev. = 12.1; 7 females). To enable parallel analyses across the two datasets, the developmental data in this dataset were divided into three age groups, to approximately match the division in Dataset 1: early childhood (n=6; ages 4-6, average = 5.4, st. dev. = 0.46), middle childhood (n=38; ages 7-9, average = 7.8, st. dev. = 0.65), and late childhood (n=23; ages 10-16 average = 11.5, st. dev. = 1.38).

A subset of the children and adults in Dataset 2 wore a total light-exclusion blindfold during the fMRI scan to reduce visual cortex responses to visual input during the scan, for a different study^102^ (children: n=14 [early = 2; middle = 7; late = 5], ages 4-16, average =8.85 , st. dev. = 2.23; adults: n=16, ages 18-59, average = 43.8, st. dev. =12.08). Blindfolded and non-blindfolded children are treated as a single group for the current study, which focuses on responses in higher-cognitive networks.

Subsets of Datasets 1 and 2 were included in several published studies^103–106,41^, whose goals differed from the current study and did not concern the development of the language network.

### 2. Language localizer task

As shown in **Figure 1**, four variants of the language localizer were used, with the control condition varying between speech played in reverse, acoustically degraded speech, and speech in an unfamiliar foreign language (see **SI-1A** for details). Participants in Dataset 2 completed a short comprehension task following each language stimuli (see **SI-1A** for task details and **SI-1B** for accuracy data). Each participant completed 6-12 blocks per condition across 1-4 scanning runs (for the early childhood group in Dataset 1, where a single run was used, the run was manually split into two equal-sized runs to enable across-runs cross-validation, as needed for the analyses; see <u>Critical individual-level neural measures</u>). Participants were excluded from the language localizer task analysis if more than 40% of the acquired volumes (across both runs) were identified as outlier volumes during preprocessing (as detailed below).

In addition to the language localizer task, most participants in Dataset 1 completed a resting state scan (described below) and most participants completed additional scanning tasks that were unrelated to the current study.

### 3. Resting state scan

The majority of participants in Dataset 1 completed a resting state scan. The participants in the early childhood group were asked to keep their eyes open and to think about anything that came to mind. A fixation cross was shown on the screen, but the children were not instructed to keep their eyes on it. The scan lasted 5 min 15 s. The participants in the middle and late childhood groups were asked to keep their eyes on the fixation cross and to let their mind wander. The scan lasted 6 min and 22 s for the middle childhood group and 5 min and 46 s for the late childhood group. For the adult group, following^71^, participants were asked to close their eyes but to stay awake and to let their mind wander; the projector was turned off, and the lights were dimmed. The scan lasted 5 min.

Participants were excluded from the resting state analysis if more than 40% of the acquired resting state volumes were identified as outlier volumes during preprocessing (as detailed below). Of the 206 and 91 adults that were included in the language localizer analyses, 189 children (ages 4 – 14, average = 10.93, st. dev. = 3.15, 97 females) and 90 adults (ages 19-45, average = 27.38, st. dev. = 5.54, 47 females) were included in the resting state analyses (n=6 early and n=14 middle participants were excluded because of excessive motion). The developmental participants included: early childhood (n=35; ages 4-6, average = 5.78, st. dev. = 0.69), middle childhood (n=47; ages 7-10, average = 9.04, st. dev. = 0.78), and late childhood (n=107; ages 12-14, average = 13.44, st. dev. = 0.68).

### 4. fMRI data acquisition

All data were collected on a 3-Tesla Siemens Prisma scanner at the Athinoula A. Martinos Imaging Center at the McGovern Institute for Brain Research at MIT.

#### Dataset 1

##### Early Childhood

Whole-head, high-resolution T1-weighted multi-echo MPRAGE structural images were acquired in 176 sagittal slices (TR = 2,530 ms, TE = 1.64 ms/3.5 ms/5.36 ms/7.22 ms, TI = 1,400 ms, flip angle = 7°, resolution = 1.00 mm isotropic). Whole-brain functional blood oxygenation level dependent (BOLD) data were acquired using an echoplanar imaging (EPI) T2*-weighted sequence in 41 transverse slices (2 mm thick) in an interleaved order (TR = 2,500 ms, TE = 30 ms, flip angle = 90°, bandwidth = 2,298 Hz/Px, echo spacing = 0.5 ms, FoV = 192 mm, phase encoding A > P direction, in-plane resolution = 3 mm × 3 mm). Resting state scans were acquired using an EPI T2*-weighted sequence in 41 sagittal slices (3 mm thick) in an interleaved order with a 10% distance factor (TR = 5200 ms, TE=30 ms, flip angle = 55°, bandwidth = 1,502 Hz/Px, FoV = 192 mm, phase encoding A > P direction, in-plane resolution = 3 mm x 3 mm). Prospective acquisition correction (PACE) was used to adjust the position of the gradients based on the participant’s motion one TR back.

##### Middle Childhood

Whole-head, high-resolution T1-weighted multi-echo MPRAGE structural images were acquired in 176 sagittal slices (TR = 2,530 ms, TE = 1.69 ms/3.55 ms/5.41 ms/7.27 ms, TI = 1,400 ms, flip angle = 7°, resolution = 1.00 mm isotropic). Whole-brain functional BOLD data were acquired using an EPI T2* weighted sequence in 32 near-axial slices (4 mm thick) in an interleaved order with a 10% distance factor (TR = 2,000 ms, TE = 30 ms, flip angle = 90°, bandwidth = 1,860 Hz/Px, echo spacing = 0.63 ms, FoV = 200 mm, phase encoding A > P direction, in-plane resolution = 2.1 mm x 2.1 mm). Resting state scans were acquired using an EPI T2*-weighted sequence in 40 sagittal slices (3 mm thick) in an interleaved order with a 10% distance factor (TR = 2,500 ms, TE = 30 ms, flip angle = 90°, bandwidth = 2,380 Hz/Px, echo spacing = 0.49 mm, FoV = 210 mm, phase encoding A > P direction, in-plane resolution = 3 mm x 3 mm). PACE was used to adjust the position of the gradients based on the participant’s motion one TR back^107^.

##### Late Childhood

Whole-head, high-resolution T1-weighted multi-echo MPRAGE structural images were collected in 320 sagittal slices (TR = 4,000 ms, TE = 1.06 ms, flip angle = 2°, resolution = 2.00 mm isotropic). Whole-brain functional BOLD data were acquired using an EPI T2*-weighted sequence in 72 near-axial slices (4 mm thick) in an interleaved order with a 10% distance factor and using GRAPPA with an acceleration factor of 2 (TR = 1,000 ms, TE = 37.2 ms, flip angle = 63°, bandwidth = 2,290 Hz/Px, echo spacing = 0.58 ms, FoV = 208 mm, phase encoding A > P direction, matrix size = 96 × 96, in-plane resolution = 2 mm x 2 mm). Resting state scans were acquired using an EPI T2*-weighted sequence in 72 axial slices (2 mm thick) in an interleaved order with a 0% distance factor (TR = 800 ms, TE = 37 ms, flip angle = 52°, bandwidth = 2,290 Hz/Px, echo spacing = 0.58 mm, FoV = 208 mm, phase encoding A > P direction, in-plane resolution = 2 mm x 2 mm).

##### Adults

Whole-head, high-resolution T1-weighted multi-echo MPRAGE structural images were collected in 179 sagittal slices (TR = 2,530 ms, TE = 3.48 ms, flip angle = 7°, resolution = 1 mm isotropic). Whole-brain functional BOLD data were acquired using an EPI T2*-weighted sequence in 31 near-axial slices (4 mm thick) in an interleaved order with a 10% distance factor and using GRAPPA with an acceleration factor of 2 (TR = 2,000 ms, TE = 30 ms, flip angle = 90°, bandwidth = 1,578 Hz/px, echo spacing = 0.72 ms, FoV = 200 mm, phase encoding A > P direction, matrix size = 96 x 96, in-plane resolution = 2.1 mm x 2.1 mm).

#### Dataset 2

Whole-head, high-resolution T1-weighted structural images were collected using one of two custom 32-channel head coils made for children(Keil et al., 2011), a 12-channel coil, or the standard Siemens 32-channel head coil in 176 interleaved sagittal slices (TR = 2,530 ms, TE= 1.64/3.5 ms/5.36 ms/7.22 ms, resolution = 1 mm isotropic). Whole-brain functional BOLD data were collected in 32 near-axial slices (3 mm thick) in an interleaved order with a 20% distance factor (TR = 2,000 ms, TE = 30 ms, flip angle = 90°, bandwidth = 2298 Hz/Px, custom pediatric coils: FoV = 192 mm; standard 32-channel head coil: FOV = 256 mm; 12-channel head coil: FOV = unavailable, phase encoding A>P direction, matrix size = 64×64, in-plane resolution = 3 mm × 3 mm). PACE was used to adjust the position of the gradients based on the participant’s motion one TR back^107^.

### 5. fMRI data preprocessing and first-level modeling

fMRI data were preprocessed and analyzed using SPM12 (release 7487), CONN EvLab module (release 19b) and custom MATLAB scripts. Each participant’s functional and structural data were converted from DICOM to NIfTI format. All functional scans were co-registered and resampled using B-spline interpolation to the first scan of the first session. Potential outlier scans were identified from the resulting subject-motion estimates as well as from BOLD signal indicators using default thresholds in the CONN pre-processing pipeline (5 st. dev. or more above the mean in global BOLD signal change or framewise displacement values above 0.9 mm). Functional and structural data were independently normalized into a common space (the Montreal Neurological Institute (MNI) template, IXI549Space) using the SPM12 unified segmentation and normalization procedure with a reference functional image computed as the mean functional image after realignment across all time points, omitting outlier scans. The output data were resampled to a common bounding box between MNI-space coordinates (−90, −126, and −72) and (90, 90, and 108), using 2 mm isotropic voxels and fourth-order spline interpolation for the functional data and 1 mm isotropic voxels and tri-linear interpolation for the structural data. Lastly, the functional data were smoothed spatially using spatial convolution with a 4 mm full-width half-maximum (FWHM) Gaussian kernel.

For the critical and control conditions of the language localizer task, effects were estimated using a general linear model (GLM) in which each experimental condition was modeled with a boxcar function convolved with the canonical hemodynamic response function (HRF) (fixation was modeled implicitly). Temporal autocorrelations in the BOLD signal timeseries were accounted for by a combination of high-pass filtering with a 128 s cutoff and whitening using an AR (0.2) model (first-order autoregressive model linearized around the coefficient a = 0.2) to approximate the observed covariance of the functional data in the context of restricted maximum likelihood (ReML) estimation. In addition to main condition effects, other model parameters in the GLM design included first-order temporal derivatives for each condition (for modeling spatial variability in the HRF delays) as well as nuisance regressors to control for the effect on the BOLD signal of slow linear drifts, subject-motion parameters, and outlier scans.

The resting state data were pre-processed using the CONN toolbox with default parameters unless stated otherwise. First, to remove noise resulting from signal fluctuations originating from non-neuronal sources (for example, cardiac or respiratory activity), the first five BOLD signal time points extracted from the white matter and cerebrospinal fluid (CSF) were regressed out of each voxel’s time course. White matter and CSF voxels were identified based on segmentation of the anatomical image^108^. Second, the residual signal was band-pass filtered at 0.008–0.09 Hz to preserve only low-frequency signal fluctuations^109^.

### 6. Functional ROI (fROI) definition

For each participant, functional regions of interest (fROIs) were defined using the Group-constrained Subject-Specific (GcSS) approach^30^. For the language network in the left hemisphere (LH), we used five parcels derived from a group-level representation of the language localizer data in 220 adult participants (independent of the adult sample in the current study) and used in much past work (e.g.,^110–112,84,113,114,9,115^, *inter alia*). These parcels include three regions in the left frontal cortex (two in the inferior frontal gyrus (LIFG and LIFGorb) and one in the middle frontal gyrus (LMFG)) and two regions in the left temporal cortex (LAntTemp and LPostTemp). Individual fROIs were defined by selecting—within each parcel—the 10% of most localizer-responsive voxels based on the *t*-values for the Language>Control contrast (see^9^ for evidence that fROIs defined in this way are similar to fROIs based on a fixed statistical significance threshold). See **SI-2C** for a sample visualization of parcels and individual-subject fROIs. We additionally defined a set of language-responsive areas in the right hemisphere (RH). Following past work (e.g.,^41^, we projected the LH parcels onto the right hemisphere and selected the 10% of most localizer-responsive voxels, as in the LH. (We chose to use parcels derived from adults in order to be able to directly compare critical neural measures between children and adults, but see **SI-2A** for evidence that parcels derived directly from the child data are similar.)

### 7. Critical individual-level neural measures of language processing

Statistical analyses were performed on a set of individual-level neural measures of language processing, including i) the magnitude of neural response, ii) the volume of activation, and iii) the strength of inter-regional functional correlations during naturalistic cognition (resting state).

#### Response magnitude

We extracted the responses (in percent BOLD signal change) from each individually defined language fROI (averaging the responses across the voxels in each fROI) to each condition (Language (Intact/Forward) and Control (Degraded/Backward/Foreign language)) relative to the fixation baseline. To ensure independence between the data used to define the fROIs and to estimate their response magnitudes, we used an across-runs cross-validation procedure (e.g.,^31^). Response magnitude was averaged across run splits, resulting in one value per participant for statistical analyses.

#### Volume of activation

Following past work (e.g.,^56,84,41^, we extracted the total number of significant voxels for each fROI above the uncorrected p<0.001 for Dataset 1 and p<0.01 for Dataset 2 threshold for Language>Control contrast. At these thresholds, most participants showed suprathreshold voxels.

#### Inter-region functional correlation

For the participants with resting state data, we extracted the BOLD signal timeseries from each individually defined language fROI using the localizer-responsive voxels as defined above. We then averaged the responses across the voxels in each fROI and computed Pearson’s moment correlation coefficients between the timeseries for each pair of fROIs (45 pairwise correlations among the 10 language fROIs, 5 fROIs in each hemisphere). These correlations were Fisher-transformed to improve normality and decrease biases in averaging^116^.

#### Lateralization Index

To determine the degree of LH-lateralization based on activation volume, we extracted the number of significant voxels (see *Volume of activation* above) for each of the five bilateral language parcels and calculated the lateralization index as follows: (LH – RH) / (LH + RH). In complementary analyses, we used the LI toolbox^57^, to calculate LI based on response magnitudes within the language parcels, using best practice adaptive thresholding and bootstrapping methods^60^ (**SI-3C**).

### 8. Statistical Analyses

We asked three research questions about the development of the language network, as described next. The analyses were identical across the two datasets, except for analyses of inter-region functional correlations, which were only performed for Dataset 1. Bayesian analyses confirmed that Dataset 1 provides substantial-to-strong evidence for testing hypotheses about age-related changes in lateralization; detailed results are provided in **SI-3B**. (All of the analyses, including model specification, are available at: https://osf.io/j582b.)

#### 1) Do children (of different ages) show adult-like topography of the left hemisphere language network?

We examined whether children show a reliable response to language relative to the control condition in the LH language network overall, and in the temporal and frontal components separately (**Table 1**). For each age group in each dataset, we fit a linear mixed-effects regression model predicting the BOLD response (estimated in each region as described in <u>Critical</u> <u>individual-level neural measures)</u> from condition (Language vs. Control) with random intercepts for participants and fROIs. For completeness, we additionally fit the same model for the RH homotope of the language network (**SI-4A**).

#### 2) Do children (of different ages) show left-hemispheric bias, and is the bias adult-like?

We tested the effects of hemisphere on the two critical neural measures volume of activation and magnitude of the critical Language> Control contrast for each age group (**Table 2**). 1) Volume of activation for the Language>Control contrast was extracted by calculating, for each participant in each dataset, the number of language-responsive voxels summed across all parcels. We additionally fit the same model separately for the temporal and frontal parcels (**SI-3A**).

Based on the examination of individual whole-brain activation maps, we chose the p<0.001 uncorrected whole-brain threshold for Dataset 1 and the p<0.01 uncorrected whole-brain threshold for Dataset 2 (the activations were generally weaker for Dataset 2). 2) Magnitude of the Language>Control contrast was extracted as described in the previous section.

Next, we asked whether the LH bias was larger in adults compared to the child groups using complementary methods that control for absolute differences in the magnitude of effects, undue influence of outliers, and potential task confounds using the lateralization index following prior work (e.g.,^56,84,41,92^. These complementary analyses included: 1) Using the volume measure defined above, we calculated a lateralization index (LI) based on the number of significantly activated voxels per hemisphere using the following formula: (LH − RH) / (LH + RH), where LH and RH represent the number of significant voxels in the left and right hemispheres, respectively. We fit a linear regression model for each dataset predicting LI using age group as a categorical predictor, followed by pairwise comparisons between each developmental group and the adult group, adjusted via Sidak’s method^117^. 2) For the magnitude-based measure, we fit a linear mixed-effects model to predict the magnitude of the Language > Control contrast using age group, hemisphere, and their interaction. 3) We used the the LI toolbox developed by ^57^ and ^60^, which allows for adaptive thresholding and uses bootstrapping methods to derive a threshold-independent magnitude-based LI and its confidence interval (see **SI-3C** for details).

#### 3) Do response magnitude and inter-regional correlations exhibit age-related changes?

To better understand developmental changes in other properties of the language regions, we evaluated the effects of age on the magnitude of activation for the Language>Control contrast. We fit linear mixed-effects models to predict magnitude from age group with participants and fROIs as random effects. We then performed planned pairwise comparisons between each child group and the adult group adjusting for multiple comparisons using Sidak’s method^117^. We repeated the analysis in children only with age as a continuous variable. Additionally, we examined the strength of inter-regional correlations within the left hemisphere language network during resting state. We fit linear regression model to predict correlation strength from age group categorically comparing each child group to the adult group with Sidak’s adjustment. As before, we repeated the analysis in children only with age as a continuous variable.

All analyses were performed in R v3.5.0 (R Core Team, 2013), using identical statistical thresholds (p < 0.05), and random effect structures (using the package lme4^118^. Significance of fixed effects in the models was tested in an ANOVA and fitted with restricted maximum likelihood (REML) using the package lmerTest^119^. Degrees of freedom were estimated using the Satterthwaite approximation^119,120^.

In addition to the analyses reported in the main text, we performed a version of the analyses where the total number of outlier volumes (see <u>fMRI data preprocessing and first-level</u> <u>modeling</u>) was used as a covariate. The results of the analyses are reported in **SI-4C**. The patterns of results were not affected by the inclusion of this motion-related covariate.

### 9. Data Availability

Source data are provided with this paper. Raw data, individual activation maps, derived analysis frames, and analysis code are available at OSF (https://osf.io/3mvpx/). A zipped **Source Data** folder contains the raw values underlying all figures and tables (means, individual data points for scatter/dot plots, and line-graph values).

## Acknowledgements and Funding Sources

We acknowledge the Athinoula A. Martinos Imaging Center at the McGovern Institute for Brain Research, MIT. For technical support during scanning, we thank Steve Shannon and Atsushi Takahashi. We are also grateful to Anna Greenwald and Elissa Newport for sharing the data reported in Olulade et al. (2020); and to the participants and their families for making this research possible. OO was supported by NIH award F32-HD100064 from NICHD. AO was supported by NIH awards T32-DC000038 and F31-DC020864 from NIDCD and the Hock E. Tan and K. Lisa Yang Center for Autism Research. RS was supported by the David and Lucile Packard Foundation (#2008-333024), NIH award R01-MH096914-05 from NIMH, and the Ellison Medical Foundation. JDEG was supported by the William and Flora Hewlett Foundation (#4429), the Walton Family Foundation, Chan Zuckerberg Initiative for the Reach Every Reader project, and the Halis Fund for Dyslexia Research at MIT. EF was supported by NIH awards DC016607 and DC016950 from NIDCD, and NS121471 from NINDS, and funds from the McGovern Institute for Brain Research and the Simons Center for the Social Brain at MIT.

## CRedIT Statement

Ola Ozernov-Palchik*: Conceptualization, Methodology, Investigation (DS 1), Data curation, Formal analysis, Visualization, Writing – original draft

Amanda M. O’Brien*: Conceptualization, Methodology, Formal analysis, Visualization, Writing – original draft

Elizabeth Lee: Data curation; Validation

Hilary Richardson: Methodology, Investigation (DS 2), Data curation, Writing – Review & Editing

Rachel Romeo: Methodology, Investigation (DS 1), Data curation, Writing – Review & Editing Moshe Poliak: Formal analysis; Writing – Review & Editing

Benjamin Lipkin: Data curation

Hannah Small: Data curation

Jimmy Capella: Investigation (DS 1), Data curation

Alfonso Nieto-Castañón: Data curation, Validation, Software

Rebecca Saxe: Methodology, Supervision, Writing – Review & Editing

John D. E. Gabrieli: Methodology, Supervision, Writing – Review & Editing

Evelina Fedorenko: Conceptualization, Methodology, Writing – original draft, Supervision, Project administration

## Notes

https://osf.io/3mvpx/files/osfstorage

### Competing Interest Statement

The authors have declared no competing interest.

### Summary of Updates

In response to reviewer feedback, we simplified the primary analyses (hemisphere by age model), removed the response-magnitude LI, added alternative LI and Bayes Factor analyses, and performed combined-dataset analyses to increase power. We also revised the text for clarity, tempered claims, restructured the Supplemental Materials, and provided disaggregated Source Data and code.

