## Supplemental Data 1 for "Precision fMRI reveals that the language network exhibits adult-like left-hemispheric lateralization by 4 years of age"

**Table of Contents**

**SI-1: Language Localizers**

- SI-1A. Detailed description of the language localizer paradigms.
- SI-1B. Behavioral performance of Dataset 2 participants on the in-scanner task.

**SI-2: The Language Network’s Topography in Children**

- SI-2A. Group-constrained Subject-Specific (GSS) analysis of child participants.
- SI-2B. Within- and between-participant similarity of activation patterns for the *Language > Control* contrast.
- SI-2C. Visualizations of the functional regions of interest in sample participants.

**SI-3: Additional Analyses Related to Language Lateralization**

- SI-3A. Volume and magnitude lateralization analysis for the temporal and frontal language areas separately.
- SI-3B. Bayesian analysis for the magnitude-based lateralization models.
- SI-3C. Lateralization Index (LI) for magnitude of response calculated using Wilke’s LI SPM toolbox.
- SI-3D. Lateralization analysis combining Datasets 1 and 2.
- SI-3E. Cross-hemispheric functional correlations in the language network.
- SI-3F. Ordered age comparisons for lateralization.

**SI-4: Additional Analyses Related to Other Aspects of the Data**

- SI-4A. Magnitude of responses to language in the RH language network across development. [parallels main Results section 1]
- SI-4B. Analyses that treat age as a continuous variable. [parallels main Results sections 2-3]
- SI-4C. Age-related changes in the magnitude of the Language > Control contrast in the LH language network controlling for motion. [parallels main Results section 3]
- SI-4D. Ordered age comparisons for the magnitude of the *Language > Control* contrast and the strength of functional correlations in the LH language network.

**SI-5: Evaluating the Reproducibility and Robustness of Olulade et al.’s (2020) study.**

- SI-5A. Summary of Olulade et al.’s study design and the critical lateralization analysis.
- SI-5B. Evaluating the *reproducibility* of Olulade et al.’s (2020) findings.
- SI-5C. Evaluating the *robustness* of Olulade et al.’s (2020) findings to preprocessing and modeling choices and measures of lateralization.

**SI-1: Language Localizers**

**SI-1A. Detailed description of the language localizer paradigms**.

***Localizer Variant 1*** (used for the early childhood group in Dataset 1): The critical condition consisted of short (15 s long) stories from the ‘*Narrative Language Measures’* component of the CUBED language assessment (Peterson & Spencer, 2012). The stories were recorded by a female native speaker of English and described events that children in this age range are likely to be familiar with (e.g., playing games, getting hurt). The control condition consisted of the same clips played in reverse (see e.g., Bedny et al., 2011 and Olson et al., 2025 for a similar approach). (A third condition, of no relevance for the current study, included different critical stimuli played to different ears.) The materials used in this variant, along with Variants 2 and 3 are available on OSF: https://osf.io/3mvpx/. Participants were instructed to listen to the clips, and a picture of a female stick figure appeared on a gray screen throughout auditory stimulation in both conditions to remind children to listen. Each participant performed one run, which consisted of 6 experimental events for each of the three conditions and 18 fixation periods (5 s each), for a total run duration of 360 s (**Table SI-1A**). Condition order was pseudo-randomized for each participant, so that no events from the same condition appeared in a row. (The runs were manually divided into two runs for analyses, as described in Methods-Section2.)

***Localizer Variant 2*** (used for the middle childhood and adult groups in Dataset 1): The critical condition consisted of short (18 s long) passages from *Alice’s Adventures in Wonderland* (Carroll, 2011). For the middle childhood group, we used the English language version, and for the adult group, which consisted of native speakers of diverse languages, we used the version in each participant’s native language (data from Malik-Moraleda, Ayyash et al., 2022). The passages were recorded by a female native speaker. For the control condition, the stimuli were acoustically degraded, as described originally in Scott et al. (2017). Briefly, the intact files were low-pass filtered at a pass-band frequency of 500 Hz. In addition, a noise track was created from each intact clip by randomizing 0.02-s long periods. To produce variations in the volume of the noise, the noise track was multiplied by the amplitude of the intact clip’s signal over time. The noise track was then low-pass filtered at a pass-band frequency of 8,000 Hz and a stop frequency of 10,000 Hz to soften the highest frequencies. The noise track and the low-pass-filtered copies of the intact files were then combined, and the level of noise was adjusted to a point that rendered the clips unintelligible. The resulting degraded clips sound like poor radio reception of speech, in which the linguistic content is not discernible. For all participants in the adult group and a subset of participants in the middle child group, a third condition was included, which consisted of passages in an unfamiliar foreign language. Because the acoustically degraded condition i) was presented to all participants, and ii) is the same as the control condition used in Localizer Variant 3 as well as being more similar to the reverse-speech control condition used in Localizer Variant 1, we chose to use this condition, instead of the foreign language condition, as the control condition (but see Malik-Moraleda, Ayyash et al., 2022 for evidence that the acoustically degraded and foreign-language conditions both work well as control conditions). Participants were instructed to listen to the clips. Participants who performed the two-condition version completed two runs, each consisting of 6 experimental events per condition and 4 fixation periods (23 s each), for a total run duration of 264 s. Participants who performed the three-condition version completed three runs, each consisting of 4 experimental events per condition and 3 fixation periods (12 s each), for a total run duration of 252 s. In both versions, condition order was palindromic and varied across runs.

***Localizer Variant 3*** (used for the late childhood group in Dataset 1): The critical condition consisted of short (18 s long) engaging excerpts from diverse sources (e.g., *The Moth* podcast, *TED* *Talks*, celebrity interviews), as detailed in Scott et al. (2017). For the control condition, the stimuli were acoustically degraded, as described in Scott et al. (2017) and identically to the procedure used for Localizer Variant 2. Participants were instructed to listen to the clips. Participants completed two runs, each consisting of 8 experimental events per condition and 5 fixation periods (14 s each), for a total run duration of 358 s. Condition order was palindromic and varied across runs.

***Localizer Variant 4*** (used for all participants in Dataset 2): The critical condition consisted of short (20 s long) engaging stories. The stories were recorded by one of three female native English speakers. The stories varied in content and fell into three conditions (Mental—stories about the characters’ mental states, Social—stories about the characters’ appearance and social relationships, and Physical—stories about physical objects and events in the world); for the purposes of the current study, we combined data from all stories given that in the other localizer variants, the materials in the critical language condition spanned diverse kinds of content, including mental, social, and physical types of events. The control condition consisted of clips in an unfamiliar foreign language. (A fifth condition, of no interest to the current study, included music clips.) The materials used in this Localizer Variant are available on OSF: https://osf.io/cbw6f/. Across conditions, each stimulus was followed by an auditorily presented question (“Does this come next?”; 1.5 s in duration), and then by another short (3 s) probe clip. The clip was followed by a 6.5 s pause during which participants were instructed to respond by pushing one of two buttons (“Yes” or “No”). Correct responses were followed by an encouraging phrase (“Way to go!”), and incorrect response by “Let’s try another!”. For the language stimuli, half of the stories were followed by the correct ending (and thus required the “Yes” response); incorrect endings were drawn randomly from all other English story conditions. For the control stimuli (unfamiliar foreign language stimuli in Hebrew, Korean, or Russian), half of the stories were followed by a clip in the same language (and thus required the “Yes” response), and the other half by a clip in another language. Participants completed four runs, each consisting of 2 experimental events per condition and three rest periods 12 s each), for a total run duration of 6.6 min. A rest period occurred at the start of each run, at the halfway point of each run (after five experimental events), and at the end of each run. Condition order was palindromic and varied across runs. (Only the stimuli themselves (20s stories in the participant’s native language or unfamiliar foreign language) were modeled as the experimental events.)

|  | **Variant 1**  (DS 1, Early Childhood) | **Variant 2**  (DS 1, Middle Childhood, Adults) | **Variant 3**  (DS 1, Late Childhood) | **Variant 4**  (DS 2, All Age Groups) |
| --- | --- | --- | --- | --- |
| **Number of experimental conditions** | 3 | 3 | 2 | 5 |
| **Condition names** | Critical: Forward; Control: Backward;  Other: Dichotic | Critical: Intact; Control: Degraded;  Other: Foreign | Critical: Intact; Control: Degraded | Critical: Mental, Social, Physical; Control: Foreign;  Other: Music |
| **Critical contrast** | Forward > Backward | Intact > Degraded | Intact > Degraded | Average of Mental,  Social, Physical > Foreign |
| **Task** | Attentive passive listening | Attentive passive listening | Attentive passive listening | “Does this come next?” task (see text) |
| **Experimental event duration** | 15 sec | 18 sec | 18 sec | 20 sec (+ task, modeled separately) |
| **Number of experimental events per run** | 18 (6 per condition) | 12 (4 per condition) | 16 (8 per condition) | 10 (2 per condition) |
| **Total number of experimental events for the critical and control conditions** | 6; 6 | 12; 12 | 16; 16 | 24; 8 |
| **Rest period duration** | 5 sec | 12 sec | 14 sec | 12 sec |
| **Number of rest periods per run** | 18 | 3 | 5 | 3 |
| **Run duration** | 360 sec | 252 sec | 358 sec | 396 sec |
| **Number of runs** | 1 (divided into 2) | 3 | 2 | 4 |

**Table SI-1A. Detailed description of the language localizer paradigms.** A summary of the design and timing parameters of the language localizer paradigms.

**SI-1B. Behavioral performance of Dataset 2 participants on the in-scanner task.**

As detailed in **SI-1A**, participants in Dataset 2 listened to 20-second story clips and, after each clip, were asked whether a 3-second probe sentence would come next in the story. Participants pressed a button to indicate "Yes" or "No," and then were provided with differentiated encouragement based on the accuracy of their response. All age groups showed above-chance performance, with the level of accuracy increasing with age, as expected: early childhood (52.37% ± 21.46%), middle childhood (75.60% ± 6.22%), late childhood (88.62% ± 5.07%), and adults (98.18% ± 0.85%). We compared participants' (N=82, one participant did not have valid in-scanner behavioral performance) response accuracies across age groups using a linear mixed-effects model, which predicted percent correct as a function of the main effects and interaction of condition and age, while accounting for individual differences with a random intercept for each participant. Pairwise comparisons indicate that adults performed significantly better than both the early group and the middle group. Additionally, late childhood participants performed significantly better than early childhood participants. No other pairwise comparisons reached significance.

| **Accuracy on the in-scanner task** | | | | |
| --- | --- | --- | --- | --- |
|  | ***B*** | ***SE*** | ***t*** | ***p*** |
| Early vs. Adult | -51.7 | 12.75 | -4.06 | **0.0008** |
| Middle vs. Adult | -26.3 | 7.49 | -3.51 | **0.0046** |
| Late vs. Adult | -12.3 | 7.57 | -1.62 | 0.38 |
| Early vs. Late | -39.4 | 12.44 | -3.17 | **0.01** |
| Middle vs. Late | -14.1 | 6.96 | -2.02 | 0.19 |
| Early vs. Middle | -25.4 | 12.40 | -2.05 | 0.18 |

**Table SI-1B. Behavioral performance of Dataset 2 participants on the in-scanner task.** Differences among the age groups in performance (accuracies) on the in-scanner task for the language localizer paradigm used for Dataset 2.

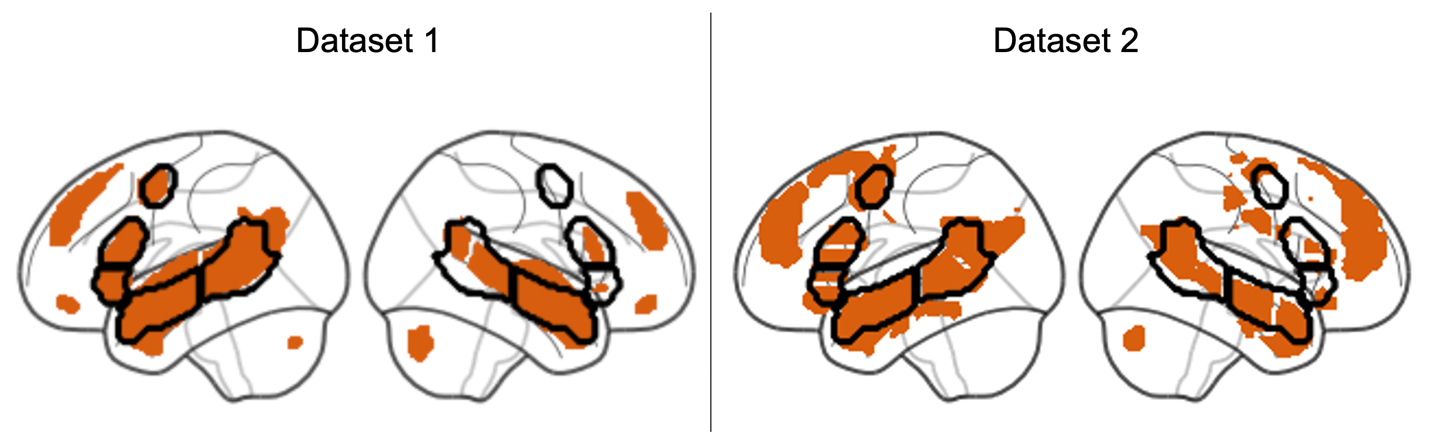
**SI-2A. Group-constrained Subject-Specific (GSS) analysis of child participants**.

**SI-2: The Language Network’s Topography in Children**

**Figure SI-2A.** **Group-constrained Subject-Specific (GSS) analysis of the child participants.** In all analyses in the current paper, we used language parcels (masks) that were derived from adult data (in prior work). This approach makes comparisons with adults (and with numerous prior studies) straightforward, but also makes an assumption that the general topography of the language activations is consistent across development. To test whether this assumption is warranted, we performed a GSS analysis (Fedorenko et al., 2010; Julian et al., 2012) on the child datasets to see if the parcels that are derived from child data are similar to those derived from adults. To perform this analysis, we followed the procedure outlined in Lipkin et al. (2022): in particular, we i) selected the top 10% of voxels across the brain for the *Language > Control* contrast from each participant, ii) binarized the resulting maps so that voxels in the top 10% set get assigned a value of 1 and the rest of the voxels – 0, iii) overlaid the binarized maps in the MNI space to derive a probabilistic overlap map, and finally, iv) parcellated the overlap map using a watershed algorithm. The parcels that resulted for each child dataset are shown in orange, and the adult parcels are overlaid in black outlines. As can be seen, the adult parcels overlap near-perfectly with the parcels derived from the child data, at least in the left hemisphere (in the right hemisphere, the child parcels look more noisy, in line with weaker responses in the right hemisphere and overall weaker responses in younger participants). It is worth noting that the parcels in the child data that are not included in the adult set (e.g., the medial frontal parcels, or the right cerebellar parcels) do exist in the adult data (see e.g., Fedorenko et al., 2010; Lipkin et al., 2022; Wolna et al., 2025), but are excluded from the adult set due to the current paper’s focus on the ‘core’ lateral frontal and lateral temporal areas.

**SI-2B. Within- and between-participant similarity of activation patterns for the *Language > Control* contrast.**

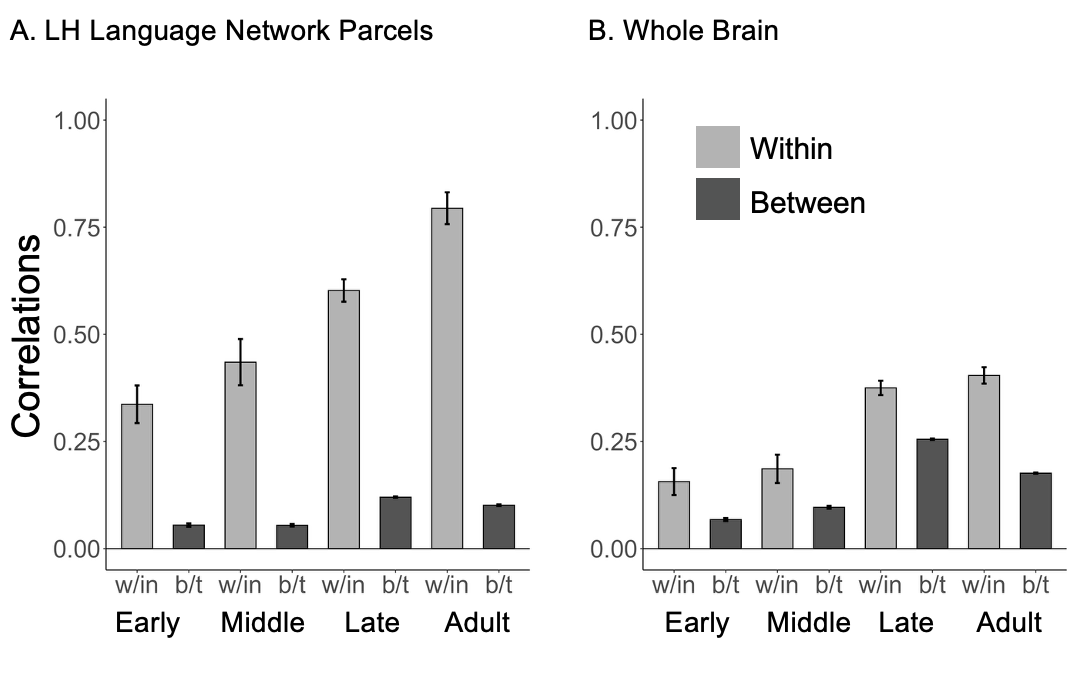

**Figure SI-2B. Within- and between-participant similarity of activation patterns for the *Language > Control* contrast for the language parcels (A) and across the brain (B).** (See **Table SI-2B** for statistical comparisons.) This analysis was performed using Dataset 1 to evaluate the stability of individual-level activation maps for the *Language > Control* contrast across runs for the different age groups (the Within, w/in bars, lighter grey), and the degree of inter-individual variability in the activation patterns (the Between, b/t bars, darker grey). **A.** The within-participant (across-run) and between-participant correlations of the activation patterns within the LH language parcels. The between-participant correlations were computed by correlating the activations of each participant with that of every other participant in that age group, and then averaging the values to derive a single value per participant. **B.** Same as in A, but not restricting the voxels to fall within the LH language parcels.

Three aspects of the results are worth highlighting. 1) Both in the LH parcels and across the brain, the within-participant correlations are substantially higher than the between-participant correlations for all age groups, which underscores the importance of identifying the language areas within individual participants, including in developmental studies. 2) The stability of the activations across runs (the Within bars) increases with age, in line with the increasing strength of the response to language (**Fig. 4** in the main text) (these two quantities typically go together; e.g., Lipkin et al., 2022). 3) The topographic similarity of a given participant to other participants (the between bars) also increases with age, albeit more modestly, which suggests that activation topographies are more idiosyncratic in younger children, which makes individual-level functional localization even more critical for developmental studies.

| **A. Age differences in within-participant correlations (indexing activation stability across runs) within the LH language network.** | | | | |
| --- | --- | --- | --- | --- |
|  | ***B*** | ***SE*** | **t** | ***p*** |
| Age = Continuous | 0.04 | 0.01 | 5.80 | **<0.001** |
| Early vs. Adult | -0.46 | 0.06 | -7.22 | **<0.001** |
| Middle vs. Adult | -0.36 | 0.06 | -6.54 | **<0.001** |
| Late vs. Adult | -0.19 | 0.05 | -4.00 | **0.001** |
| **B. Age differences in within-participant correlations across the brain.** | | | | |
|  | ***B*** | ***SE*** | **t** | ***p*** |
| Age = Continuous | 0.03 | 0.00 | 6.69 | **<0.001** |
| Early vs. Adult | -0.25 | 0.04 | -6.70 | **<0.001** |
| Middle vs. Adult | -0.22 | 0.03 | -6.56 | **<0.001** |
| Late vs. Adult | -0.03 | 0.03 | -1.04 | 0.88 |
| **C. Age differences in between-participant correlations (indexing inter-individual topographic variability) within the LH language network.** | | | | |
|  | ***B*** | ***SE*** | **t statistic** | ***p*** |
| Age = Continuous | -0.01 | 0.00 | -10.39 | **<0.001** |
| Early vs. Adult | -0.05 | 0.01 | -6.31 | **<0.001** |
| Middle vs. Adult | -0.05 | 0.01 | -7.04 | **<0.001** |
| Late vs. Adult | -0.02 | 0.01 | -3.42 | **0.004** |
| **D. Age differences in between-participant correlations across the brain.** | | | | |
|  | ***B*** | ***SE*** | **t** | ***p*** |
| Age = Continuous | 0.03 | 0.00 | 15.30 | **<0.001** |
| Early vs. Adult | -0.11 | 0.12 | -9.15 | **<0.001** |
| Middle vs. Adult | -0.08 | 0.01 | -7.56 | **<0.001** |
| Late vs. Adult | 0.08 | 0.01 | 8.45 | **<0.001** |

**Table SI-2B. Age differences in the within- (A, B) and between-participant (C, D) similarity of activation patterns for the *Language > Control* contrast for the language parcels (A, C) and across the brain (B, D).**

For all child participants in Dataset 1, we fit a linear regression model predicting the correlation coefficients from age as a continuous variable (the Age = Continuous rows). In addition, we fit a linear regression model on all participants in Dataset 1 using age group as a categorical predictor, followed by pairwise comparisons between the adult group and each child group, adjusted using Sidak’s method (Methods-Section8).

**SI-2C. Visualizations of the functional regions of interest in sample participants.**

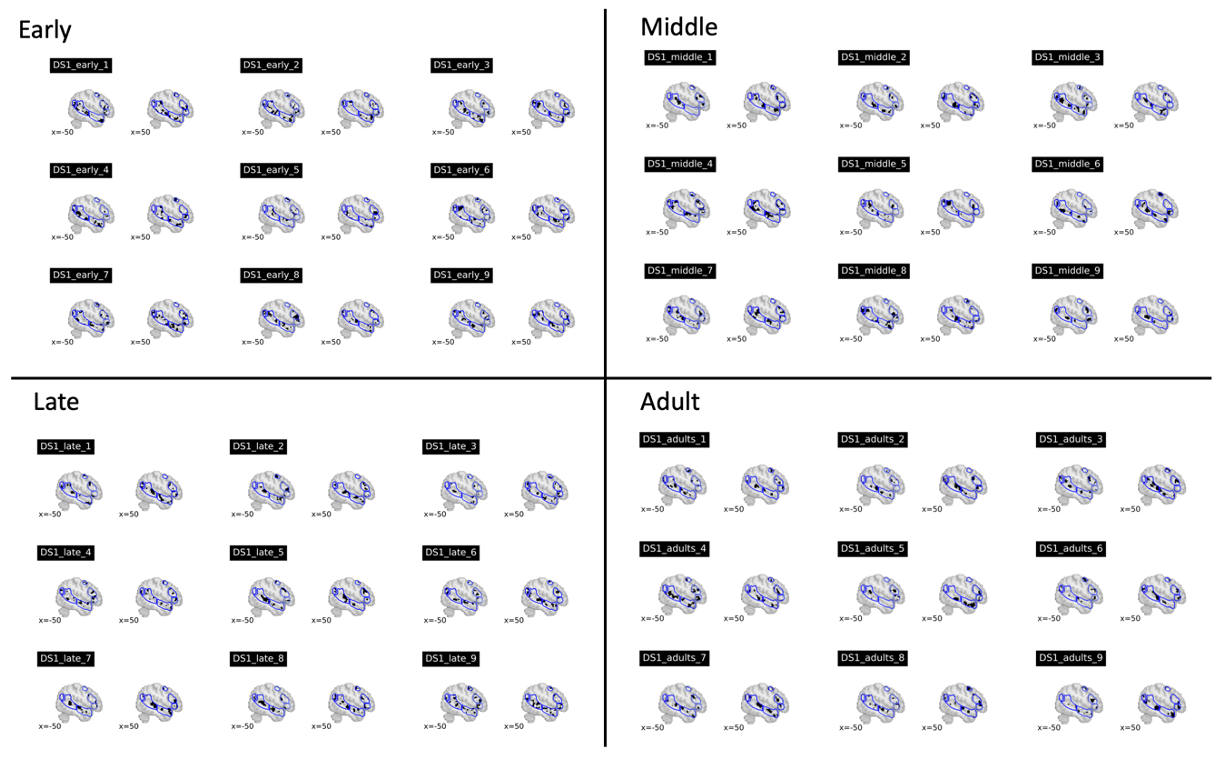

**Figure SI-2C**. **Visualization of the functional regions of interest in sample participants from each age group in Dataset 1.** The fROIs are defined as the 10% of most language-responsive voxels within the language parcels (see Methods-Section6). All the individual maps are available at: <https://osf.io/3mvpx/files/osfstorage>.

**SI-3: Additional Analyses Related to Language Lateralization**

**SI3-A. Volume and magnitude lateralization analysis for the temporal and frontal language areas separately.**

| **A. Volume-based lateralization index (LI)** [complementary to main Table 3A] | | | | | | | | | | | | |
| --- | --- | --- | --- | --- | --- | --- | --- | --- | --- | --- | --- | --- |
|  | | Frontal | | | | | Temporal | | | | | |
|  | | *b* | *SE* | *t* | | *p* | *b* | *SE* | | *t* | | *p* |
| DS 1 | Early vs. Adult | 0.14 | 0.11 | 1.28 | | 0.492 | 0.21 | 0.07 | | 2.88 | | **0.01** |
|  | Middle vs. Adult | -0.08 | 0.10 | -0.81 | | 0.804 | -0.09 | 0.07 | | -1.35 | | 0.44 |
|  | Late vs. Adult | 0.00 | 0.08 | -0.02 | | 1.000 | 0.01 | 0.05 | | 0.12 | | 1.00 |
| DS 2 | Early vs. Adult | -0.24 | 0.28 | -0.86 | | 0.393 | -0.07 | 0.21 | | -0.33 | | 0.78 |
|  | Middle vs. Adult | -0.11 | 0.16 | -0.70 | | 0.488 | -0.05 | 0.13 | | -0.40 | | 0.87 |
|  | Late vs. Adult | -0.05 | 0.18 | -0.26 | | 0.799 | -0.07 | 0.14 | | -0.48 | | 0.99 |
| **B. Magnitude-based lateralization** [complementary to main Table 3A] | | | | | | | | | | | | |
|  | | Frontal | | | | | Temporal | | | | | |
|  | | *b* | *SE* | | *t* | *p* | *b* | *SE* | *t* | | *p* | |
| DS 1 | Early vs. Adult | 0.23 | 0.18 | 1.26 | | 0.502 | 0.32 | 0.14 | | 2.27 | | 0.29 |
|  | Middle vs. Adult | -0.32 | 0.17 | -1.97 | | 0.144 | -0.01 | 0.18 | | -0.04 | | 1.00 |
|  | Late vs. Adult | 0.03 | 0.14 | 0.23 | | 0.994 | 0.47 | 0.11 | | 4.36 | | **0.01** |
| DS 2 | Early vs. Adult | -0.35 | 0.30 | -1.15 | | 0.581 | -0.16 | 0.20 | | -0.79 | | 0.89 |
|  | Middle vs. Adult | 0.30 | 0.19 | -1.57 | | 0.321 | -0.12 | 0.13 | | -0.95 | | 0.82 |
|  | Late vs. Adult | 0.04 | 0.21 | 0.20 | | 0.996 | 0.13 | 0.14 | | 0.91 | | 0.84 |

**Table SI-3A. No evidence for an age-related increase in the LH bias for language.** **A**. **Volume-based LI analysis.** We fit a linear regression model predicting LI (averaged across the three frontal fROIs (Frontal) and across the two temporal fROIs (Temporal)) from age group, followed by pairwise comparisons between the adult group and each child group, adjusted using Sidak's method. **B. Magnitude-based lateralization analysis.** We fit a linear mixed-effect regression model predicting the size of the *Language > Control* contrast in the Frontal or in the Temporal component from age group, hemisphere, and their interaction, with random intercepts and slopes for hemisphere within participants and fROIs, followed by pairwise comparisons, as in A. (Note that the two significant effects go in the opposite direction of the one predicted by the hypothesis we are evaluating(Lenneberg, 1967).)

**SI-3B. Bayesian analysis for the magnitude-based lateralization models**

**Overview.** Given that the primary finding in the current study is the *lack of support* for increased lateralization with age, we sought to confirm this finding with a Bayesian analysis, which allow for quantification of evidence in favor of the null hypothesis (Kass & Raftery, 1995; Nicenboim et al., 2025). Bayes Factors represent the ratio of marginal likelihoods between two competing models, essentially quantifying how much more likely the observed data are under one model compared to another. For instance, a Bayes Factor of 10 indicates that the data are ten times more likely under the model in the numerator than under the comparison model, whereas a Bayes Factor of 1/10 indicates the reverse. A Bayes Factor of 1 means the data are equally likely under both models. When comparing nested models that differ only by the inclusion of a single predictor (e.g., the effect of age on lateralization), the Bayes Factor can directly quantify evidence for or against that predictor's contribution. In our case, we evaluated the null hypothesis that lateralization does not change with age against the alternative hypothesis that it does. Following established conventions, Bayes Factors between 0.312 and 3.2 indicate anecdotal (weak) evidence, values exceeding 3.2 (or below 1/3.2) represent substantial evidence, and values exceeding 10 (or below 1/10) constitute strong evidence (Kass & Raftery, 1995).

**Model fitting.** For each dataset, we fit Bayesian models using the *brms* package in R (Bürkner, 2018) that mirror the frequentist models in the main analysis. Specifically, we predicted the difference in response magnitude for the Language > Control contrast from hemisphere, age, and their interaction, with random intercepts and slopes for hemisphere within participants and fROIs.

We fit two types of models for each dataset, differing in how age group was coded. The first type of model imposed a monotonic constraint on age effects (Bürkner & Charpentier, 2020). Under this constraint—consistent with the primary hypothesis of increased lateralization with age—the main effect of age on activation could only change in one direction (e.g., increase with age). Although this constraint allows for non-linear changes (e.g., lateralization increasing from early to middle childhood, then plateauing), it excludes more complex developmental trajectories. For instance, it would not permit lateralization to be high in early childhood, decrease during middle childhood, and then return to high levels in adulthood. In the second type of model, we evaluated each child group against the adult group, consistent with our focus on examining whether lateralization is already adult-like in the youngest childhood group, without imposing directional constraints.

All models were fit with 4 chains of 20,000 (4,000 warmup) iterations and a step size of 0.999. For all models, the R-hat values for all parameters were 1, indicating good model convergence. All models converged with no divergent transitions except for 4 models fit on Dataset 2, which had up to 21 divergent transitions after warmup. We do not consider this issue worrisome because (1) these divergent transitions constitute (at most) .00105 of all iterations, (2) no abnormalities were apparent upon visual inspection, and (3) the estimates are consistent across the various models (including ones without divergent transitions; Betancourt, 2017). In total, we fit 10 models per dataset (5 using the monotonicity constraint and 5 without), varying in how the priors are specified, as detailed next.

**Priors.** Bayesian modeling explicitly incorporates prior beliefs about parameters, unlike frequentist modeling where such assumptions remain implicit (Wald, 1950; Jaynes, 1968). We chose priors centered at 0, representing a neutral starting point that allows for effects in either direction without favoring one over the other. The width of these priors reflects our beliefs about the magnitude of potential developmental effects on lateralization. A Normal(0, sd) prior reflects the belief that there is a roughly 95% probability that the effect falls within ±1.96*sd. For example, a Normal(0, .2) prior over the interaction in a model with a monotonic age constraint implies a belief that, with a ~95% probability, the average difference in lateralization between adjacent age groups is ±.392. Across all four age groups, this allows for total change up to 1.176 from early childhood to adulthood.

Each group of models (with vs. without the monotonic age constraint) consisted of one null model (no hemisphere by age interaction(s)) and four models with the interaction term(s), varying in how the prior was specified: liberal (very large expected effect), moderate (large effect consistent with previous literature), conservative (smaller than literature estimates), and extra-conservative (effect too small to be of theoretical importance and/or too small to estimate given existing datasets) (**Table SI-3B-i**). The varying widths of these priors reflect different degrees of skepticism about effect magnitude, informed by the previous literature showing moderate to large developmental differences in lateralization. In the non-null models with a monotonic constraint, a single interaction term is used, and in the non-null models without the monotonic constraint, three interaction terms are used: one per age group comparison (adults vs. early, adults vs. middle, and adults vs. late). Within each non-monotonic model, the three interaction priors were identical. For all random effects, we used brms default priors.

| **Term** | **Prior** | **Interpretation** |
| --- | --- | --- |
| Intercept | Normal(0, 2) | Overall activation difference (language > control) |
| Residuals | Normal(0, 2) | Model residual error |
| Hemisphere | Normal(0, 1) | Overall hemispheric difference in activation |
| Age | Normal(0, 1) | Overall age-related change in activation |
| Hemisphere:Age (liberal) | Normal(0, .5) | Very large changes in lateralization with age (up to ~3.0 total change from early childhood to adulthood) |
| Hemisphere:Age (moderate) | Normal(0, .2) | Moderate changes consistent with literature (up to ~1.2 total change) |
| Hemisphere:Age (conservative) | Normal(0, .1) | Small changes in lateralization (up to ~0.6 total change) |
| Hemisphere:Age (extra conservative) | Normal(0, .05) | Very small changes (up to ~0.3 total change) |

**Table SI-3B-i. Summary of Priors on fixed effects.** The priors for the intercept, residuals, hemisphere, and age were the same across all models. The prior for the interaction varied across models, expressing different prior beliefs about the potential size of the interaction effect. In the monotonic models, the prior is over the average change in lateralization per age level (early childhood → middle childhood → late childhood → adulthood). In the categorical models, the prior is over the difference between adulthood and each of the childhood groups.

**Results.**

***No evidence for monotonic change in lateralization with age.*** In Dataset 1, models testing for monotonic age effects provided **substantial-to-strong evidence against** this hypothesis across all prior specifications (BF01 = 12.67, 4.37, 2.45, and 1.84 for liberal, moderate, conservative, and extra-conservative priors, respectively; **Table SI-3B-ii**). Note that, similar to frequentist 95% Confidence Intervals, a 95% Credible Interval (CrI) is considered as evidence for a substantial effect in either direction if it does not contain 0 (Nicenboim et al., 2025). The extra-conservative model was the most favorable to the presence of an interaction (although still pointing against it). It showed substantial evidence for left-hemispheric lateralization in early childhood (right hemisphere showing lower activation: estimate = -.55, 95% CrI = [-.94, -.14]) and substantial evidence for increased left hemisphere activation with age (estimate = .29, 95% CrI = [.19, .39]). However, critically, there was no substantial evidence for a *change in lateralization* *with age* (estimate = .01, 95% CrI = [-.07, .08]), suggesting that a monotonic change in lateralization across development is either non-existent or vanishingly small.

Likely due to the very small sample size of some of the subgroups, Dataset 2 provided only anecdotal evidence for either hypothesis across all models. The most favorable model (conservative priors; BF10 = 1.66) showed no substantial evidence for lateralization in early childhood (estimate = -.29, 95% CrI = [-.66, .11]), age effects (estimate = .12, 95% CrI = [-.04, .28]), or the critical interaction (estimate = -.08, 95% CrI = [-.18, .03]). The inconclusive Bayes Factors and credible intervals containing zero suggest this dataset is weakly informative.

***No evidence for stronger lateralization in adults compared to children.*** In Dataset 1, models that treated age group as a categorical variable, comparing adults to each child group, showed substantial evidence against the null hypothesis in moderate and conservative prior models (BF10 = 6.41 and 4.30, respectively). However, all interaction estimates had credible intervals containing zero: early childhood (estimate = -.17, 95% CrI = [-.41, .08]), middle childhood (estimate = .16, 95% CrI = [-.07, .39]), and late childhood (estimate = -.15, 95% CrI = [-.35, .05]). In line with the analyses reported in the main text, the early childhood group showed greater lateralization than adults, which is the opposite of the hypothesis we are evaluating. Dataset 2 provided inconclusive evidence across all models (BF01 = 2.99, 0.66, 0.65, and 0.78), with all interaction estimates containing zero: early childhood (estimate = .03, 95% CrI = [-.15, .21]), middle childhood (estimate = .08, 95% CrI = [-.08, .23]), and late childhood (estimate = -.07, 95% CrI = [-.23, .09]).

|  | **Monotonic BF_01_ (BF_10_)** | | **Categorical BF_01_ (BF_10_)** | |
| --- | --- | --- | --- | --- |
|  | DS1 | DS2 | DS1 | DS2 |
| Liberal (large effects) | **12.67 (.08)** | 1.52 (.61) | .58 (1.73) | 2.99 (.33) |
| Moderate (medium effects) | **4.37 (.23)** | .78 (1.29) | .16 (6.41) | .66 (1.52) |
| Conservative (small effects) | 2.45 (.41) | .60 (1.66) | .23 (4.30) | .65 (1.54) |
| Extra Conservative (very small effects) | 1.84 (.54) | .61 (1.64) | .43 (2.35) | .78 (1.29) |

**Table SI-3B-ii. Bayes Factors for monotonic and categorical models testing age effects on lateralization.** BF₀₁ values indicate evidence in favor of the null hypothesis (no age effect on lateralization), while BF₁₀ values (in parentheses) indicate evidence in favor of the alternative hypothesis. Values >3.2 indicate substantial evidence, and values >10 indicate strong evidence. Liberal through extra-conservative refer to different prior specifications reflecting varying beliefs about effect magnitude (see Priors).

**Summary.**

The Bayesian analyses show support against the monotonic increase in lateralization with age, with Dataset 1 providing substantial-to-strong evidence in favor of the null hypothesis (BF01 ranging from 1.84 to 12.67). For models that compared adults to each child age group, although some models showed evidence against the null hypothesis, all parameter estimates had credible intervals containing zero, indicating no substantial developmental differences. These findings suggest that lateralization either does not change across development or changes so little that it cannot be reliably detected even with a sample size as large as 297 participants, and thus might be of limited theoretical importance.

**SI-3C. Lateralization Index (LI) for magnitude of response calculated using Wilke’s LI SPM toolbox.**

In complementary analyses to our primary magnitude analysis, we used the LI toolbox (Wilke & Lidzba, 2006; Wilke & Lidzba, 2007) to calculate LI based on response magnitude within the language parcels using best practice adaptive thresholding and bootstrapping methods with 1,000 iterations. This approach reduces the impact of outliers, specific thresholds, and data sparsity on the LI calculation by generating a threshold-independent, weighted bootstrapped mean LI using the formula LI = (L-R)/(L+R), where L and R represent the sum of activation values across ROIs in the left and right hemispheres, respectively.

| **Magnitude-based lateralization index (LI)** [complementary to main Table 3] | | | | | |
| --- | --- | --- | --- | --- | --- |
|  |  | *b* | *SE* | *t* | *p* |
| DS 1 | Early vs. Adult | 0.002 | 0.07 | 0.03 | 1.000 |
|  | Middle vs. Adult | -0.16 | 0.06 | -2.46 | 0.04 |
|  | Late vs. Adult | 0.09 | 0.53 | 1.67 | 0.26 |
| DS 2 | Early vs. Adult | -0.14 | 0.18 | -0.81 | 0.81 |
|  | Middle vs. Adult | -0.25 | 0.11 | -2.27 | 0.08 |
|  | Late vs. Adult | -0.08 | 0.12 | -0.66 | 0.88 |

**Table SI-3C. No evidence for an age-related increase in the LH bias for language using Wilke’s magnitude-based LI measure (Wilke & Lidzba, 2006).** We fit a linear regression model predicting LI (averaged across fROIs) from age group, followed by pairwise comparisons between the adult group and each child group, adjusted using Sidak's method.

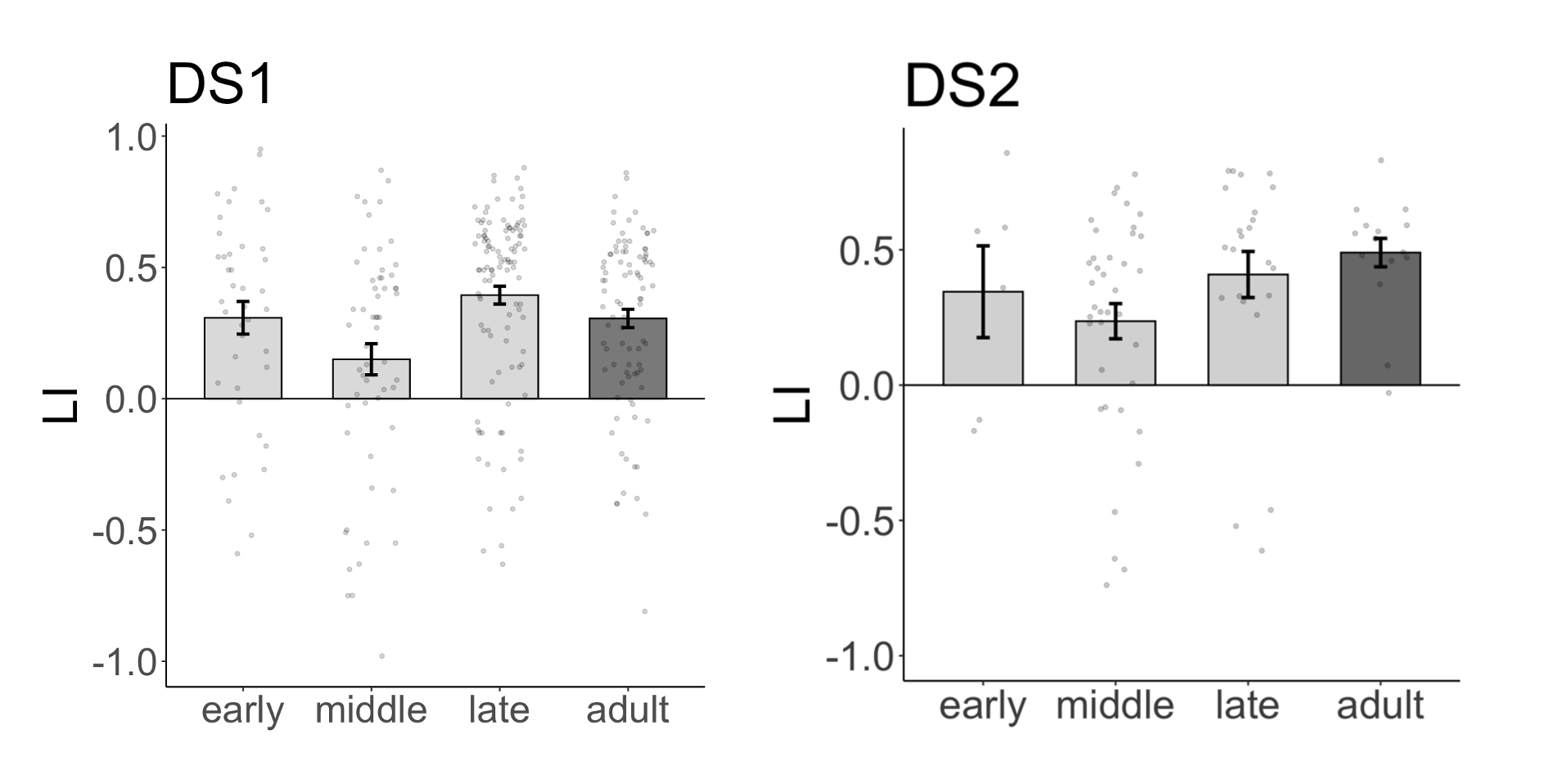

**Figure SI-3C. Developmental changes in lateralization with LI estimated using Wilke’s magnitude-based LI measure (Wilke & Lidzba, 2006).** For all bars, dots correspond to individual participants, and error bars indicate standard errors of the mean by participant (see **Table SI-3C** for statistical results). The critical finding reported in the main text—that the degree of language lateralization is not lower in children compared to adults—is present in both datasets for this additional measure of lateralization, evidencing robustness to analytic choices.

**SI-3D: Lateralization analysis combining Datasets 1 and 2.**

For the lateralization analyses using our two main measures (volume-based LI and a magnitude-based lateralization measure), we conducted an analysis combining data across the two datasets. Both models for this combined dataset were sufficiently powered: the power of volume-based LI model was estimated at 91.73% (Cohen’s f² = 0.04), and the power of the magnitude-based LI model was estimated at 99.92% (Cohen’s f² = 0.125). Thus, both models exceed the conventional 80% threshold, indicating that our analyses are well‐powered to detect even differences between age groups. Crucially, the results for these analyses parallel those obtained when each dataset was analyzed separately: no evidence for an age-related increase in lateralization.

| **A. Volume-based lateralization index (LI) using the combined dataset**  [parallel to main Table 3A] | | | | | |
| --- | --- | --- | --- | --- | --- |
|  | | *b* | *SE* | *t* | *p* |
| DS 1 and DS 2 | Early vs. Adult | 0.16 | 0.07 | 2.27 | 0.07 |
|  | Middle vs. Adult | -0.03 | 0.06 | -0.61 | 0.91 |
|  | Late vs. Adult | -0.01 | 0.05 | -0.13 | 1.00 |
| **B. Magnitude-based lateralization using the combined dataset**  [parallel to main Table 3B] | | | | | |
|  |  | *b* | *SE* | *z* | *p* |
| DS 1 and DS 2 | Early vs. Adult | 0.20 | 0.15 | 1.35 | 0.44 |
|  | Middle vs. Adult | -0.20 | 0.12 | -1.66 | 0.26 |
|  | Late vs. Adult | 0.18 | 0.11 | 1.69 | 0.25 |

**Table SI-3D. No evidence for an age-related increase in the LH bias for language, even when combining the data across the two datasets.** **A. Volume-based LI analysis.** As in the main analyses, we fit a linear regression model predicting LI (averaged across fROIs) from age group, followed by pairwise comparisons between the adult group and each child group, adjusted using Sidak’s method. **B. Magnitude-based lateralization analysis.** As in the main analyses, we fit a linear mixed-effect regression model predicting the size of the *Language > Control* contrast from age group, hemisphere, and their interaction, with random intercepts and slopes for hemisphere within participants and fROIs, followed by pairwise comparisons, as in A.

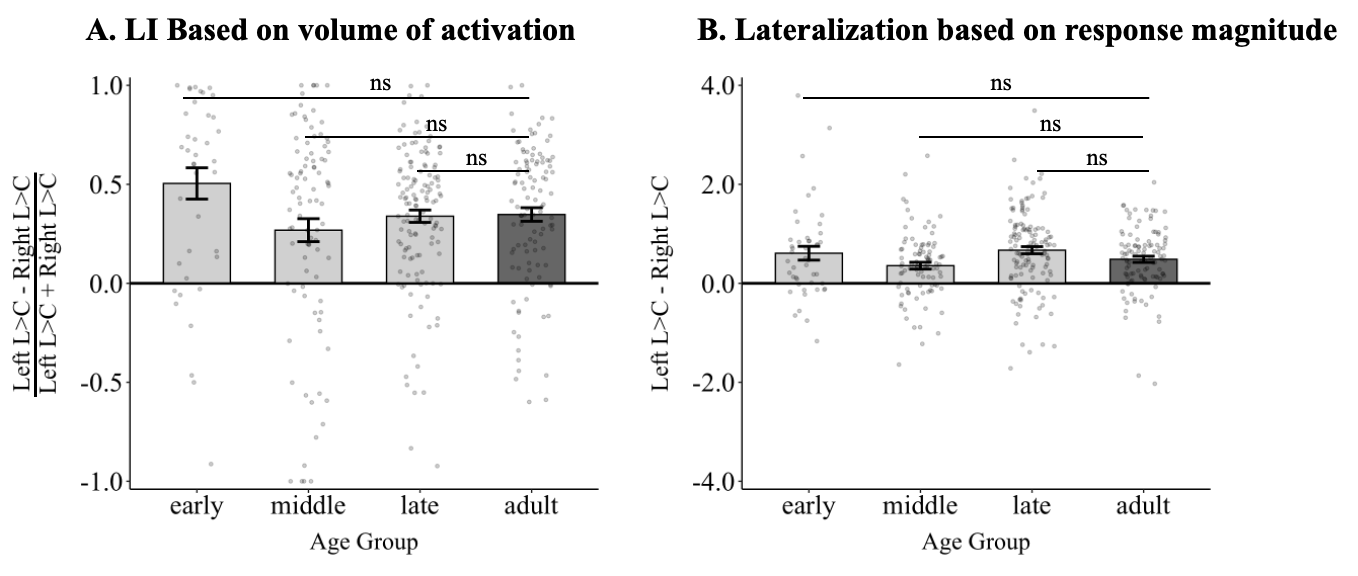

**Figure SI-3D. Developmental changes in lateralization. A.** Volume-based LI measure for participants combined across Datasets 1 and 2 in each child group (light grey bars) and the adult group (dark grey bar). Here and in B, no significant differences were found between children and adults for any child group. **B.** Magnitude-based lateralization measure. For all bars, dots correspond to individual participants, and error bars indicate standard errors of the mean by participant. All comparisons between child groups and adults were non-significant (see **Table SI-3D** for details).

**SI-3E. Cross-hemispheric functional correlations in the language network.**

|  |  | *b* | *SE* | *t* | *p* |
| --- | --- | --- | --- | --- | --- |
| DS 1 | Early vs. Adult | -0.07 | 0.03 | -2.05 | 0.12 |
|  | Middle vs. Adult | -0.22 | 0.03 | -7.04 | **<0.001** |
|  | Late vs. Adult | -0.03 | 0.02 | -1.09 | 0.62 |

**Table SI-3E.** **No evidence for an age-related change in the strength of cross-hemispheric functional correlations in the language network.** We computed functional correlations for pairs of homotopic language regions in Dataset 1 and averaged the five values to obtain a single value per participant. We then fit a linear mixed-effects regression model predicting cross-hemispheric correlation strength from age group, with random intercepts for participants and ROIs, followed by pairwise comparisons between the adult group and each child group, adjusted using Sidak's method.

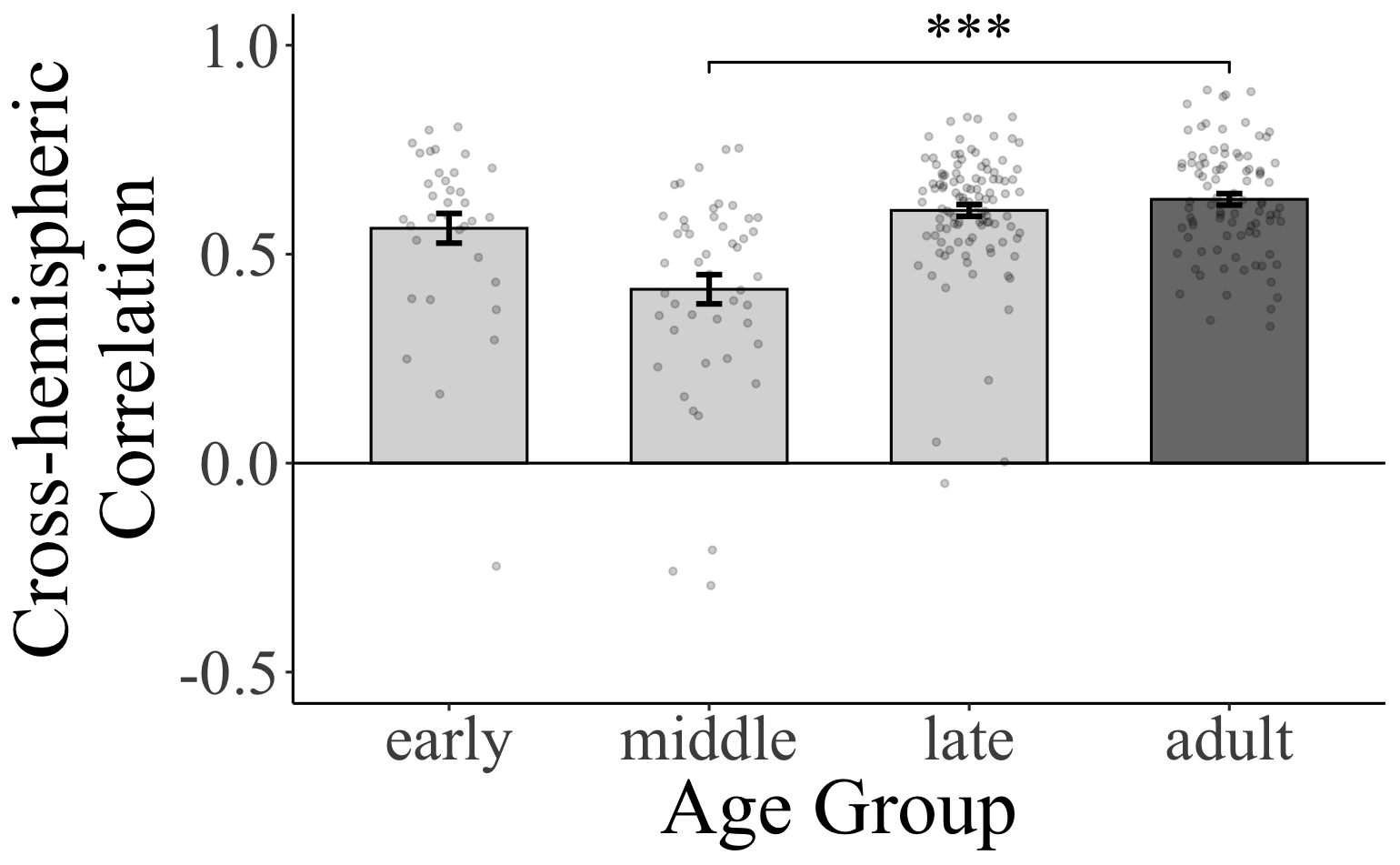

**Figure SI-3E. Cross-hemispheric functional correlations.** Functional correlations for pairs of homotopic language regions in Datasets 1 in each child group (light grey bars) and the adult group (dark grey bar). Significance: ***=*p*<0.001 (see **Table SI-3E** for details).

**SI-3F. Ordered age comparisons for lateralization.**

| **A. Volume-based lateralization index (LI)** | | | | | |
| --- | --- | --- | --- | --- | --- |
|  | | *b* | *SE* | *t* | *p* |
| DS 1 | Early vs. Late | 0.17 | 0.07 | 2.56 | **0.03** |
|  | Middle vs. Late | -0.09 | 0.06 | -1.55 | 0.33 |
|  | Early vs. Middle | 0.26 | 0.08 | 3.48 | **0.002** |
| DS 2 | Early vs. Late | -0.09 | 0.17 | -0.52 | 0.94 |
|  | Middle vs. Late | -0.02 | 0.10 | -0.24 | 0.99 |
|  | Early vs. Middle | -0.07 | 0.17 | -0.41 | 0.97 |
| **B. Magnitude-based lateralization** | | | | | |
|  | | *b* | *SE* | t | *p* |
| DS 1 | Early vs. Late (LH vs. RH) | 0.06 | 0.18 | 0.34 | 0.981 |
|  | Middle vs. Late (LH vs. RH) | -0.40 | 0.16 | -2.51 | **0.04** |
|  | Early vs. Middle (LH vs. RH) | 0.46 | 0.20 | 2.31 | 0.07 |
| DS 2 | Early vs. Late (LH vs. RH) | -0.35 | 0.26 | -1.34 | 0.46 |
|  | Middle vs. Late (LH vs. RH) | -0.30 | 0.15 | -2.00 | 0.14 |
|  | Early vs. Middle (LH vs. RH) | -0.05 | 0.25 | -0.19 | 1.00 |

**Table SI-3F. No evidence for an age-related increase in the LH bias for language.** **A**. **Volume-based LI analysis.** We fit a linear regression model predicting LI (averaged across fROIs) from age group, followed by ordered pairwise comparisons between the adult group and each child group, adjusted using Sidak’s method. **B. Magnitude-based lateralization analysis.** We fit a linear mixed-effect regression model predicting the size of the *Language > Control* contrast from age group, hemisphere, and their interaction, with random intercepts and slopes for hemisphere within participants and fROIs, followed by pairwise comparisons, as in A.

**SI-4: Additional Analyses Related to Other Aspects of the Data**

**SI-4A: Magnitude of responses to language in the RH language network across development.** [parallels main Results section 1]

| **Magnitude of responses to the *Language > Control* contrast in the RH language network**  [parallel to main Table 1] | | | | | | | | | | | | | |
| --- | --- | --- | --- | --- | --- | --- | --- | --- | --- | --- | --- | --- | --- |
|  | | RH language network | | | | RH temporal language areas | | | | RH frontal language areas | | | |
|  |  | *B* | *SE* | *t* | *p* | *B* | *SE* | *t* | *p* | *B* | *SE* | *t* | *p* |
| DS 1 | Early | 0.16 | 0.08 | 1.95 | 0.05 | 0.55 | 0.11 | 5.05 | **<0.001** | -0.1 | 0.11 | -0.88 | 0.378 |
|  | Middle | 0.82 | 0.08 | 9.96 | **<0.001** | 1.34 | 0.11 | 11.85 | **<0.001** | 0.46 | 0.10 | 5.21 | **<0.001** |
|  | Late | 1.13 | 0.05 | 21.14 | **<0.001** | 1.82 | 0.08 | 22.69 | **<0.001** | 0.67 | 0.05 | 12.15 | **<0.001** |
|  | Adult | 1.06 | 0.06 | 17.95 | **<0.001** | 1.52 | 0.09 | 17.68 | **<0.001** | 0.76 | 0.07 | 11.57 | **<0.001** |
| DS 2 | Early | 0.19 | 0.12 | 1.64 | 0.11 | 0.41 | 0.22 | 1.91 | 0.07 | 0.04 | 0.12 | 0.37 | 0.715 |
|  | Middle | 0.44 | 0.06 | 7.45 | **<0.001** | 0.65 | 0.08 | 7.85 | **<0.001** | 0.3 | 0.08 | 3.84 | **<0.001** |
|  | Late | 0.43 | 0.07 | 6.58 | **<0.001** | 0.67 | 0.1 | 6.82 | **<0.001** | 0.27 | 0.09 | 3.21 | **0.002** |
|  | Adult | 0.35 | 0.06 | 5.61 | **<0.001** | 0.50 | 0.08 | 5.98 | **<0.001** | 0.24 | 0.08 | 3.07 | **0.003** |

**Table SI-4A. Magnitude of response to language in the RH language network as a whole, and in the temporal and frontal components separately, across development**. For each age group in each dataset, we fit a linear mixed-effects regression model predicting the magnitude of the BOLD response in each region from condition (Language vs. Control) with random intercepts for participants and fROIs. The responses are estimated in data independent from the data used to define the language fROIs. The *p*-values for the temporal and frontal areas are uncorrected, but all significant effects survive a Bonferroni correction for two comparisons.

**SI-4B: Analyses that treat age as a continuous variable.** [parallels main Results sections 2-3]

|  | *B* | *SE* | *t* | *p* |
| --- | --- | --- | --- | --- |
| **A. Volume-based lateralization index (LI)**  [parallel to main Table 3A] | | | | |
| DS 1 | -0.01 | 0.02 | -0.51 | 0.61 |
| DS 2 | -0.02 | 0.03 | -0.55 | 0.59 |
| **B. Magnitude-based lateralization**  [parallel to main Table 3B] | | | | |
| DS 1 | -0.02 | 0.02 | -0.95 | 0.35 |
| DS 2 | -0.03 | 0.03 | -0.96 | 0.34 |
| **C. Magnitude of the *Language > Control* contrast in the LH language network**  [parallel to main Table 4A] | | | | |
| DS 1 | 0.13 | 0.02 | 5.66 | **< 0.001** |
| DS 2 | 0.065 | 0.034 | 1.91 | 0.06 |
| **D. Inter-regional functional correlation strength in the LH language network**  [parallel to main Table 4B] | | | | |
| DS 1 | 0.02 | 0.00 | 8.20 | **< 0.001** |

**Table SI-4B. Effects of age (as a continuous variable) on neural measures of language processing in children.** For each analysis, we fit regression models predicting neural responses from age, with random intercepts included where appropriate. Adults were excluded. **A.** Linear regression model predicting the volume-based lateralization index (LI) from age. **B.** Linear mixed-effects regression model predicting effect magnitude from hemisphere and age, with random intercepts for participants and fROIs. **C.** Linear regression model predicting the magnitude of the Language > Control contrast in the LH language network. **D.** Linear regression model predicting the strength of inter-regional functional correlations during resting state (averaged across the 10 LH fROI pairs; available only for Dataset 1).

**SI-4C: Age-related changes in the magnitude of the *Language > Control* contrast in the LH language network controlling for motion.** [parallels main Results section 3]

| **A. Magnitude of the *Language > Control* contrast in the LH language network, controlling for motion** | | | | | |
| --- | --- | --- | --- | --- | --- |
|  |  | *b* | SE | t | *p* |
| DS1 | Early vs. Adult | -0.60 | 0.17 | -3.58 | **0.001** |
|  | Middle vs. Adult | -0.02 | 0.18 | -0.11 | 0.99 |
|  | Late vs. Adult | 0.37 | 0.14 | 2.69 | **0.02** |
| DS2 | Early vs. Adult | 0.41 | 0.29 | -1.43 | 0.40 |
|  | Middle vs. Adult | -0.01 | 0.20 | -0.06 | 1.00 |
|  | Late vs. Adult | 0.21 | 0.20 | 1.06 | 0.65 |
| **B. Inter-regional functional correlation strength in the LH language network, controlling for motion** | | | | | |
|  | Early vs. Adult | -0.11 | 0.02 | -4.16 | **< 0.001** |
|  | Middle vs. Adult | -0.09 | 0.02 | -3.74 | **0.002** |
|  | Late vs. Adult | 0.02 | 0.02 | 1.23 | 0.44 |

**Table SI-4C.** **Evidence for** **age-related changes in language response magnitude and functional connectivity in the LH language network after controlling for motion. A. Magnitude of the *Language> Control* contrast analysis.** We fit a linear mixed-effects regression model predicting response magnitude of the LH language network from age group, with random intercepts for participants and fROIs, and critically, including motion as a covariate. We then performed pairwise comparisons between the adult group and each child group, adjusted using Sidak’s method. **B. Inter-regional functional correlation strength analysis.** We fit a linear regression model predicting inter-regional correlations (averaged across the 10 fROI pairs for the LH network) from age group, including a motion covariate, followed by pairwise comparisons, as in A. It is also worth noting that age was not a significant predictor of motion in DS1 (F(2,188)=3.27, p=0.07) but a significant predictor in DS2 (F(3,79)=7.61, p<0.001).

**SI-4D. Ordered age comparisons for the magnitude of the *Language > Control* contrast and the strength of functional correlations in the LH language network.**

| **A. Age differences in the magnitude of the Language>Control contrast** | | | | | |
| --- | --- | --- | --- | --- | --- |
|  |  | *b* | *SE* | *t* | *p* |
| DS1 | Early vs. Late | -0.90 | 0.19 | -4.70 | **< 0.001** |
|  | Middle vs. Late | -0.74 | 0.17 | -4.33 | **< 0.001** |
|  | Early vs. Middle | -0.16 | 0.21 | -0.75 | 0.84 |
| DS2 | Early vs. Middle | -0.30 | 0.27 | -1.10 | 0.62 |
|  | Middle vs. Late | -0.29 | 0.16 | -1.82 | 0.20 |
|  | Early vs. Late | -0.59 | 0.28 | -2.11 | 0.11 |
| **B. Age differences in the strength of inter-regional correlations** | | | | | |
|  |  | *b* | *SE* | *t* | *p* |
| DS1 | Early vs. Middle | -0.03 | 0.03 | -1.38 | 0.43 |
|  | Middle vs. Late | -0.12 | 0.02 | -6.14 | **< 0.001** |
|  | Early vs. Late | -0.16 | 0.02 | -7.1 | **< 0.001** |

**Table SI-4D. Evidence for age-related increases in language response magnitude and functional connectivity in the LH language network.** We conducted linear mixed-effects analyses predicting response magnitude (A) and inter-regional correlation strength (B) from Age with random intercepts for participants and fROIs. We then performed pairwise comparisons of the adjusted means for Age using Sidak’ method for multiple comparison adjustments.

**SI-5: Evaluating the Reproducibility and Robustness of Olulade et al.’s (2020) study.**

**SI-5A. Summary of Olulade et al.’s study design and the critical lateralization analysis.**

Olulade et al. (2020) examined neural responses during a language task in a group of n=53 participants across four age groups (n=39 children and 14 adults; the age range for the children was similar to the range in the current study). Each participant performed a single run of a 2-condition experiment. In the *critical* (forward speech) condition, participants listened to short sentences and were asked to press a button at the end to decide whether the sentence was factually correct. In the *control* (backward speech) condition, participants listened to the same sentences played backwards and were asked to press a button to decide whether a beep was present at the end of the sound file. No fixation condition was included.

The key claims in Olulade et al. are that i) the language system is more bilateral in children and becomes increasingly left-lateralized with age, and ii) the change in the degree of lateralization is due to a decrease in the activity of the RH language areas with age. The following analysis was performed by Olulade and colleagues to evaluate these claims:

- Individual-level activation maps were thresholded at p<0.001 uncorrected at the whole-brain level and evaluated against two large anatomical right-hemisphere masks (ROIs). One mask covered the inferior frontal gyrus, and the other covered lateral temporal and parietal cortex.
- For each age group and for each ROI: if a participant had a cluster of size=269 voxels or larger, they were assigned a value of 1, and otherwise they were assigned a value of 0.
- For each age group and for each ROI, the values were summed, divided by the number of participants in that group, and multiplied by 100% to convert from proportions to percentages.

Thus the final measure for each age group and each right-hemisphere ROI corresponds the percentage of participants who had a cluster of size 269 voxels or larger. By this measure, Olulade et al. found that in the youngest age group, almost every participant showed language activation within the RH ROIs, but in the older age groups, the percentage of participants with RH activation decreased. These results appear in Figure 4 in Olulade et al.’s paper (repeated in the top row in **Figure SI-5B** below).

This result appears inconsistent with the results we report for our data, where we do not see a change in language lateralization as a function of age in either of the two datasets (or when combining the data across datasets; **SI-3D**): in our data, even the youngest children show left-lateralized responses, and the strength of lateralization is similar to that observed in the adult group. However, the comparison with Olulade et al.’s results is challenging given the differences between the studies in the paradigms, the preprocessing and modeling pipelines, and the measure used to assess lateralization. To facilitate the comparison, we performed two analyses, as described below. First, in Section **SI-5B**, we show that Olulade et al.’s results are reproducible by implementing their preprocessing and modeling pipeline and using their measure of lateralization. And second, in Section **SI-5C**, we show that when using our pipeline and more standard measures of lateralization, the results are not consistent with Olulade et al.’s critical claims and instead align with the findings in the current study. We then elaborate on the possible source of the discrepancy.

**SI-5B. Evaluating the *reproducibility* of Olulade et al.’s (2020) findings.**

Data: Olulade et al.’s data.

Preprocessing + analysis: Olulade et al.’s pipeline with one change in how outlier timepoints were modeled.

Lateralization measure: Olulade et al.’s measure (percentage of participants with significant activation in the large anatomical RH frontal and temporo-parietal masks).

We ran the data from Olulade et al.’s study through a preprocessing and modeling pipeline that was designed to be as similar as possible to the one used in their study. We thank Elissa Newport and Anna Greenwald for providing us with 3D nifti files, realignment parameter (rp) files, bad scan regressor files, segmentation images for WM and GM, and ROI masks. The only change we made to the modeling pipeline was to use a GLM regressor for *each* outlier scan rather than a single regressor for all outlier scans as in Olulade et al.

*[Motivation for the change in how the outliers are modeled*: Using a single regressor for all outlier scans is problematic. If a) the number of outliers varies between conditions (what is commonly referred to as “task-related motion”), and b) outliers are not modeled correctly, the residual/uncorrected outlier effects can show up as artifactual task effects (inflating both the effect sizes and the associated statistics). Because motion effects are rather large compared to typical task effects, they can easily dominate the results. Indeed, in Olulade et al.’s data, participants (and especially younger kids) moved much more during the control (reverse speech) condition compared to the critical (forward speech) condition. As a result, we wanted to ensure that their results held when the outliers were modeled correctly.]

To additionally evaluate the robustness of the results to the choice of the number of voxels above which a participant is considered to have RH activation, we report the results for four thresholds: at least 1 significant voxel, at least 10 significant voxels, at least 100 significant voxels, and at least 200 significant voxels.

We find that the results reported in Olulade et al. (included here in the top row in **Figure SI-5B** below, for ease of comparison) are reproducible when the outliers are modeled by including a regressor for each outlier scan and are robust to the choice of the threshold, showing the same downward trend in the percentage of participants with significant RH activation with age in both the frontal and temporo-parietal ROIs. However, the effects are substantially reduced in size: for example, for the frontal ROI, the difference reported in Olulade et al. between the youngest group and adults is ~75%, but in our re-analysis, the differences are on the order of 20-60% across thresholds, which suggests that improper modeling of motion outliers may have contributed to the results reported in Olulade al.’s paper.

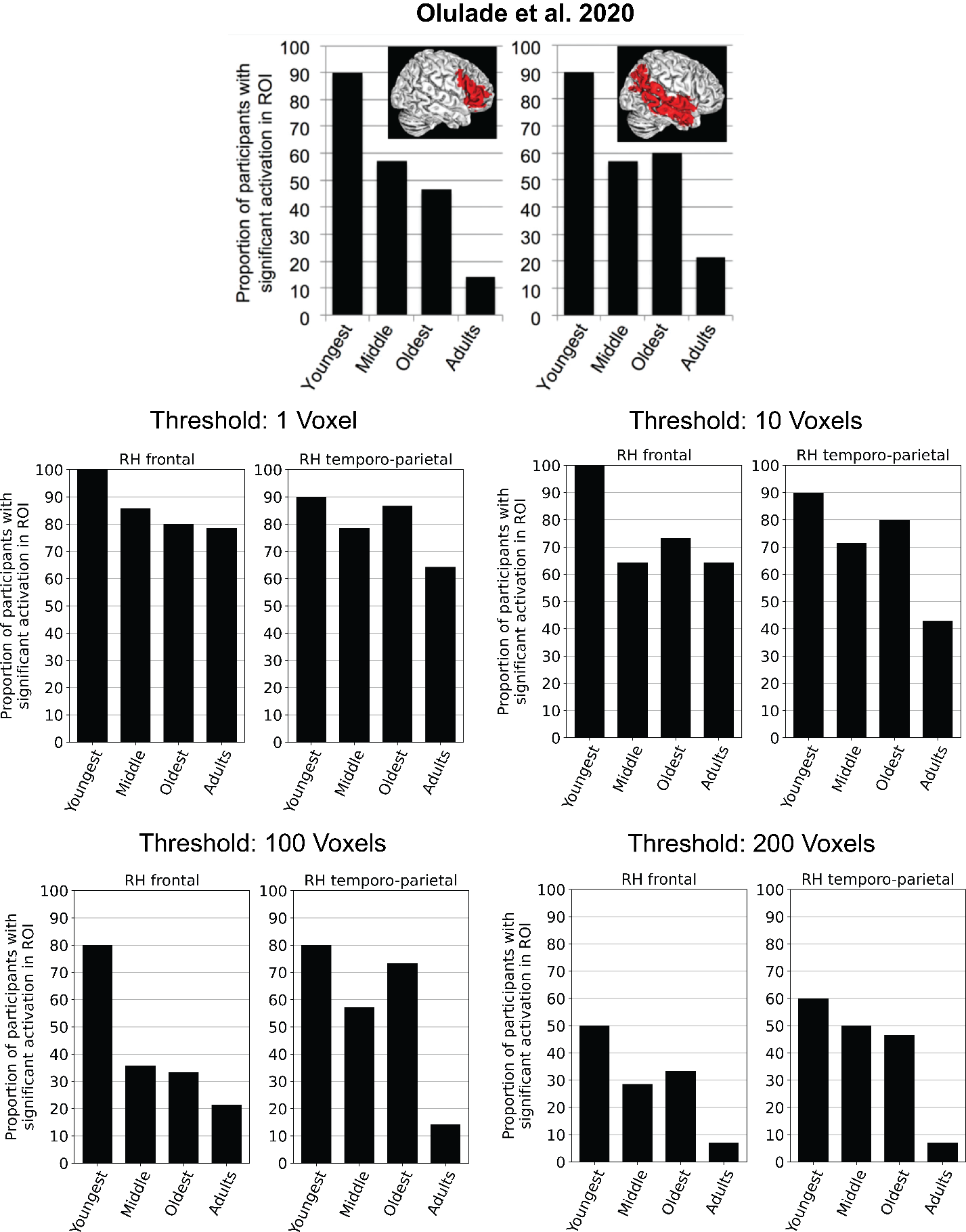

**Figure SI-5B:** A comparison between Olulade et al.’s (2020) results as reported in the paper (top row) vs. a re-analysis of their data (middle and bottom rows). The data are re-analyzed in a slightly modified modeling pipeline (as discussed above) and the lateralization effects are computed across several thresholds of how many voxels are needed to count the participant as having RH activation (1 voxel, 10 voxels, 100 voxels, and 200 voxels). The pattern reported by Olulade et al. is reproducible and robust to the choice of threshold, although the effects are reduced in size.

**SI-5C. Evaluating the *robustness* of Olulade et al.’s (2020) findings to preprocessing and modeling choices and measures of lateralization.**

Data: Olulade et al.’s and our data.

Preprocessing + analysis: The pipeline used in the current study.

Lateralization measure: A measure used in the current study (volume of activation index) and the number of LH vs. RH langu­­­­­­age-responsive voxels within the language masks; see **Methods-Section6**).

We ran the data from Olulade et al.’s study through the preprocessing and modeling pipeline that was used in the current study. The key differences between the two pipelines are summarized in Table **SI-5C**.

|  | ***Olulade et al. 2020 pipeline*** | ***Current pipeline*** |
| --- | --- | --- |
| Outlier regressors | Single regressor for all outlier scans | A regressor per outlier scan |
| Smoothing of functional data | 8mm FWHM kernel | 4mm FWHM kernel |
| Temporal response function | Canonical hemodynamic response function (HRF) | Canonical hemodynamic response function (HRF) and first-order temporal derivative |
| Global scan regressor | Included | Not included |
| Task design blocks | The first experimental block of the run is specified as starting at the 2^nd^ instead of the 1^st^ TR (1 TR = 3 sec difference). | The first experimental block of the run is specified as starting at the 1^st^ TR. |

**Table SI-5C.** Key differences between Olulade et al.’s (2020) vs. our preprocessing and modeling pipelines.

Before diving into the results, it is worth noting that Olulade et al. motivate their lateralization measure as follows (bolding is ours to highlight the critical bits of the argument):

*“The vast majority of studies to date have investigated language lateralization using a laterality index (LI) comparing LH and RH activation. While the precise method for quantifying LH and RH activation and computing the LI differs across studies (40, 41), the LI in general compares the difference between LH and RH activation with the total activation. LIs near 0 indicate bilateral and equal activation in the two hemispheres, whereas positive LIs indicate left lateralization. This measure allows quantification of lateralization regardless of absolute activation levels, which may be influenced by age or other factors that may differ over age, such as stimulus complexity, effort, and task difficulty (17, 42–45). However, because the LI is a difference score,* ***potentially important information may be lost****. An increase in LI with age could reflect a decrease of RH activation, an increase of LH activation, or both; the extent to which the LH and the RH are each involved in language processing is not directly reflected in LI scores. […] For these reasons, the present fMRI study focuses on* ***activation patterns in each hemisphere*** *rather than on lateralization per se […].”*

However, Olulade et al. do not report the amount of LH and RH activation (in magnitude or spatial extent) separately; they only report the *percentage of participants with RH activation* (above some fixed number of voxels; see **SI-5B** above). Below, we report the standard LI measure as well as volumes of LH and RH activation.

The voxel count data (bottom row in **Figure SI-5C**) reveal that **Olulade et al.’s data are not consistent with the key empirical claim in their paper** (i.e., that RH activation in the language areas decreases with age). In particular, in their dataset, the number of language-responsive voxels in the RH remains relatively constant across the four age groups (similar to our Dataset 2), but the number of language-responsive voxels in the LH increases. This pattern leads to a slight age-related increase in the lateralization index scores, from ~0.4 in the early child group to ~0.6 in adults (top row in **Figure SI-5C**). Note, however, that all four groups—including the youngest children—already show *strongly left-lateralized language activations* (all LI measures are positive, and the number of LH language-responsive voxels is larger than the number of RH language-responsive voxels), similar to what we observe in both of our datasets (second and third columns in **Figure SI-5C**).

So, what is the **possible source of the discrepancy** between Olulade et al.’s reported results (reproduced here qualitatively in **SI-5B** above) and our re-analysis of their data? As briefly mentioned in the main text, the difference plausibly has to do with i) Olulade et al.’s reliance on a paradigm that conflates linguistic and general cognitive processing, and ii) their use of masks for voxel-count analyses that encompass both language and domain-general areas. Let us unpack these points.

***First***, our study used an extensively validated language localizer paradigm (Fedorenko et al., 2010; see Fedorenko et al., 2024 for a review). This paradigm robustly differentiates the language network from the nearby domain-general Multiple Demand (MD) network (Duncan, 2010, 2013)—a bilateral network of frontal, inferior temporal, and parietal areas whose different components lie adjacent to the frontal and temporal language areas (Fedorenko et al., 2012; Fedorenko & Blank, 2020; Braga et al., 2020; DiNicola et al., 2024). The MD network is strongly sensitive to task demands across domains (e.g., Duncan & Owen, 2000; Fedorenko et al., 2013; Hugdahl et al., 2015; Shashidhara et al., 2020), and language tasks where linguistic processing is accompanied by general task demands engage the MD network in addition to the language network (Diachek, Blank, Siegelman et al., 2020).

The language task used in Olulade et al. study conflates linguistic and general task demands: the critical condition requires linguistic processing but it is also *more cognitively demanding* than the control condition. As a result, this contrast will engage both the language network and the MD network. Moreover, task demands vary across the age groups (see the fourth row in Table 1 in Olulade et al., which shows a gradual increase in the accuracy of responses across the four age groups; although the accuracies are not broken down by condition, the differences most likely stem from the critical condition given that the task in the control condition is very easy: to press a button if a beep is heard). Given these age-related differences in task difficulty, the reliance on the MD network likely also varies across age groups. In particular, younger children plausibly engage the MD network to a greater extent than older children and adults, which would result in overall more bilateral activations given that the MD network is bilateral. As the task becomes easier with age, the reliance on the MD network is likely reduced, and the language network becomes dominant, which results in more left-lateralized activations (given that the language network is typically left-lateralized).

And **second**, Olulade et al. used large anatomical masks as their ROI masks. These masks encompass parts of both the language network and the MD network (see e.g., Fedorenko & Blank, 2020; Braga et al, 2020). Given the use of a paradigm that likely engages both networks, these ROI masks capture effects that arise in both of these networks, making it impossible to unambiguously attribute the effects to one or the other network.

Future studies should either use validated localizers that have been established to isolate the language-processing mechanisms from those that support cognitive demands associated with task performance, or—if a novel task is used that potentially conflates linguistic and general cognitive processing—they should design the paradigm in a way such that the task component can be modeled separately from the language-processing component of the task (as in Dataset 2 in the current study), and/or include a localizer for the MD network to help rule out the contribution of this network if the research questions focus on the language network. Note also that the language and the MD networks can be robustly recovered at the individual participant level from the voxel-wise patterns of functional correlations in both naturalistic-cognition data (e.g., resting state data; e.g., Braga et al., 2020) and task data (e.g., Du et al., 2025; Shain & Fedorenko, 2025). So for those skeptical of functional localizers, this data-driven approach can be used.

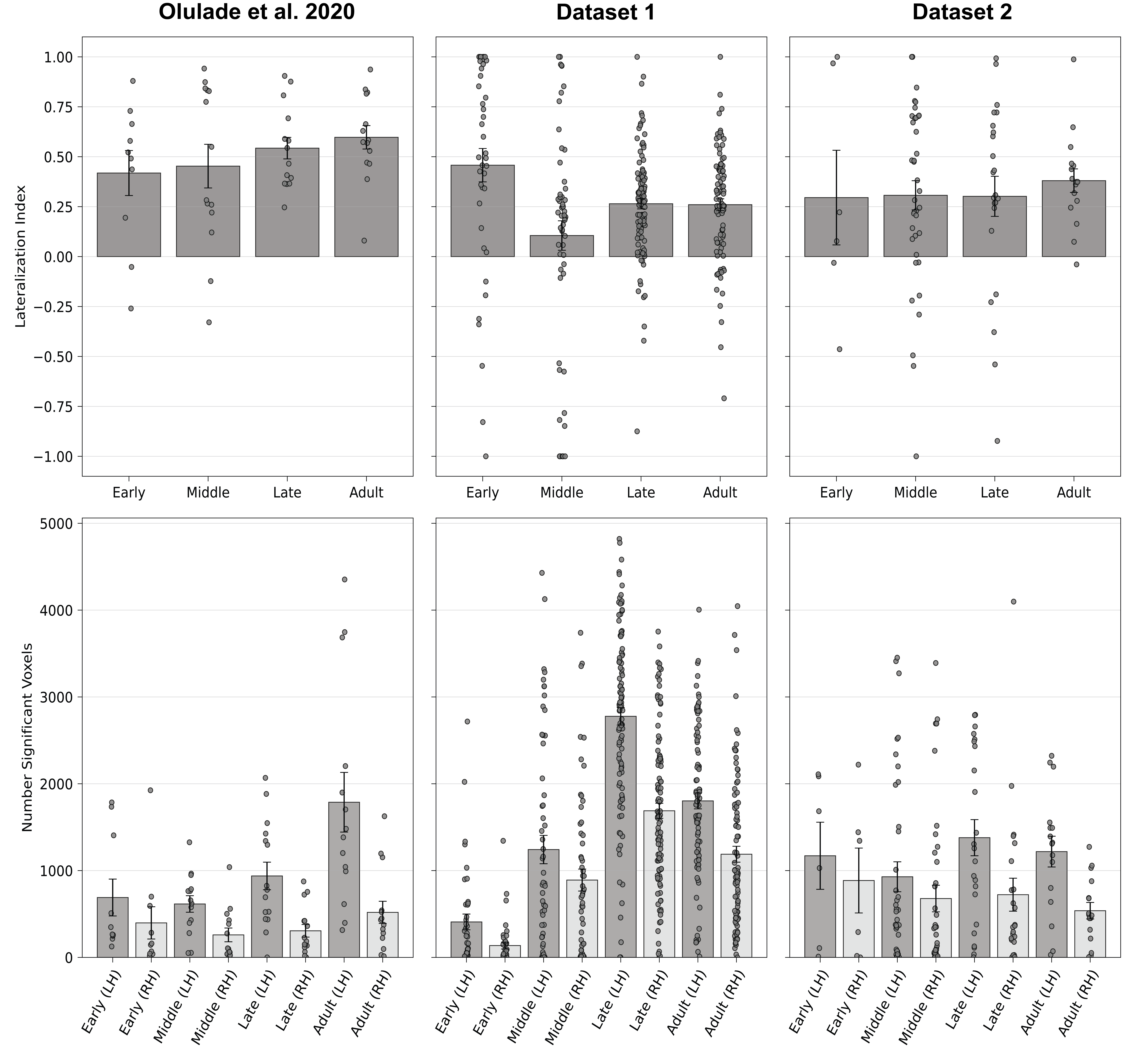

**Figure SI-5C:** A comparison between Olulade et al.’s (2020) data (left column) and our data from Dataset 1 (middle column) and Dataset 2 (right column) when all data are analyzed through the same preprocessing and modeling pipeline (used in the current study) and the lateralization effects are examined with a standard LI measure. The top row shows volume-based LI by age group, and the bottom row shows activation volumes broken down by hemisphere and by age group.
